# Glycyl-tRNA sequestration is a unifying mechanism underlying *GARS1*-associated peripheral neuropathy

**DOI:** 10.1101/2024.10.11.617948

**Authors:** Natalia Mora, Erik F.J. Slot, Vanessa Lewandowski, Maria P. Menafra, Moushami Mallik, Pascal van Lith, Céline Sijlmans, Nick van Bakel, Zoya Ignatova, Erik Storkebaum

## Abstract

Dominantly inherited mutations in eight cytosolic aminoacyl-tRNA synthetase genes cause hereditary motor and sensory neuropathy, characterized by degeneration of peripheral motor and sensory axons. We previously identified a pathogenic gain-of-toxic function mechanism underlying peripheral neuropathy (PN) caused by heterozygous mutations in the *GARS1* gene, encoding glycyl-tRNA synthetase (GlyRS). Specifically, PN-mutant GlyRS variants sequester tRNA^Gly^, which depletes the cellular tRNA^Gly^ pool, leading to insufficient glycyl-tRNA^Gly^ available to the ribosome and consequently ribosome stalling at glycine codons. Given that GlyRS functions as a homodimer, a subset of PN-GlyRS mutations might alternatively cause peripheral neuropathy through a dominant negative loss-of-function mechanism. To explore this possibility, we here generated three novel PN-GlyRS *Drosophila* models expressing human PN-GlyRS (hGlyRS) variants that do not alter the overall GlyRS protein charge (S211F and H418R) or the single reported PN-GlyRS variant that renders the GlyRS protein charge more negative (K456Q). High-level expression of hGlyRS-K456Q did not induce peripheral neuropathy and the K456Q variant does not affect aminoacylation activity, suggesting that K456Q is not a pathogenic mutation. Expression of hGlyRS-S211F or hGlyRS-H418R in *Drosophila* did induce peripheral neuropathy and *de novo* protein synthesis defects. Genetic and biochemical evidence indicates that these phenotypes were attributable to tRNA^Gly^ sequestration rather than a dominant negative mechanism. Our data identify tRNA^Gly^ sequestration as a unifying pathogenic mechanism underlying PN-GlyRS. Thus, elevating tRNA^Gly^ levels may constitute a therapeutic approach for all PN-GlyRS patients, irrespective of their disease-causing mutation.

## INTRODUCTION

Dominantly inherited mutations in eight distinct cytosolic <u>a</u>mino<u>a</u>cyl-t<u>R</u>NA <u>s</u>ynthetase (aaRS) genes give rise to Charcot-Marie-Tooth (CMT) peripheral neuropathy (1). AaRSs are the enzymes that covalently link amino acids to the 3’ end of their cognate tRNA, a reaction referred to as aminoacylation, representing the essential first step of protein biosynthesis (2–4). Heterozygous mutations in *GARS1*, the gene encoding <u>gly</u>cyl-t<u>R</u>NA <u>s</u>ynthetase (GlyRS), give rise to an axonal form of CMT, type 2D (CMT2D) (1,5). CMT is characterized by selective degeneration of peripheral motor and sensory axons, leading to progressive muscle weakness and wasting, steppage gait, sensory dysfunction, and often foot deformities (6,7). In addition to CMT2D, heterozygous *GARS1* mutations have been linked to related peripheral neuropathies characterized by selective motor axonal degeneration and lacking sensory involvement, including distal hereditary motor neuropathy type V (dHMN-V) (1,5) and infantile spinal muscular atrophy (iSMA) (8,9). Here, we refer to these *GARS1*-associated <u>p</u>eripheral <u>n</u>europathies as PN-GlyRS.

We have previously shown that six CMT-causing mutations in GlyRS or <u>tyr</u>osyl-t<u>R</u>NA <u>s</u>ynthetase (TyrRS) all inhibit protein synthesis in *Drosophila* motor and sensory neurons, uncovering the inhibition of global protein synthesis as a unifying pathogenic mechanism underlying CMT-aaRS (10). Consistent with this notion, inhibition of protein synthesis was observed in motor neurons of two PN-GlyRS mouse models (11). More recently, we identified the detailed molecular mechanism underlying PN-GlyRS: PN-GlyRS variants are still able to bind tRNA^Gly^ but fail to release it or only release at a very slow pace. This sequestration of tRNA^Gly^ by PN-GlyRS variants depletes the cellular pool of free tRNA^Gly^ available for aminoacylation by the wild type (WT) GlyRS protein in heterozygous PN-GlyRS patients. This results in insufficient production and supply of aminoacylated glycyl-tRNA^Gly^ to the ribosome, which triggers ribosome stalling at glycine codons (12). Ribosome stalling in turn induces the integrated stress response (ISR) through activation of the eIF2α kinase GCN2 (13–15), which in PN-GlyRS and CMT-TyrRS mouse models occurs selectively in motor and sensory neurons, the affected cell types in CMT (11). Consistent with this mechanism, transgenic overexpression of tRNA^Gly^ replenishes the cellular tRNA^Gly^ pool and rescues defective protein synthesis, prevents ISR activation, and rescues peripheral neuropathy phenotypes in *Drosophila* and mouse PN-GlyRS models (12).

Here, we sought to evaluate whether all PN-GlyRS mutations cause disease through tRNA^Gly^ sequestration, what can be classified as a gain-of-toxic-function mechanism, or alternatively, whether some PN-GlyRS mutations cause disease through partial loss of aminoacylation activity. Indeed, similar to tRNA^Gly^ sequestration, a substantial reduction of tRNA^Gly^ aminoacylation activity may also lead to insufficient cellular levels of aminoacylated glycyl-tRNA^Gly^ and consequently ribosome stalling at glycine codons and GCN2-mediated ISR activation (Figure 1A). Thus, although the primary molecular defect would be different, namely tRNA^Gly^ sequestration versus reduced aminoacylation activity, the downstream molecular pathway that triggers peripheral neuropathy (ribosome stalling and ISR activation) would be the same. Nevertheless, from a therapeutic perspective, the question whether some of the PN-associated GlyRS mutations may act through partial loss of aminoacylation activity is of key importance. Indeed, for tRNA^Gly^ sequestration, elevating cellular tRNA^Gly^ levels would constitute the optimal therapeutic approach, whereas for mutations resulting in partial loss of GlyRS aminoacylation activity, increasing the level of WT GlyRS protein would be the appropriate therapeutic strategy (Figure 1A).

**Figure 1:**
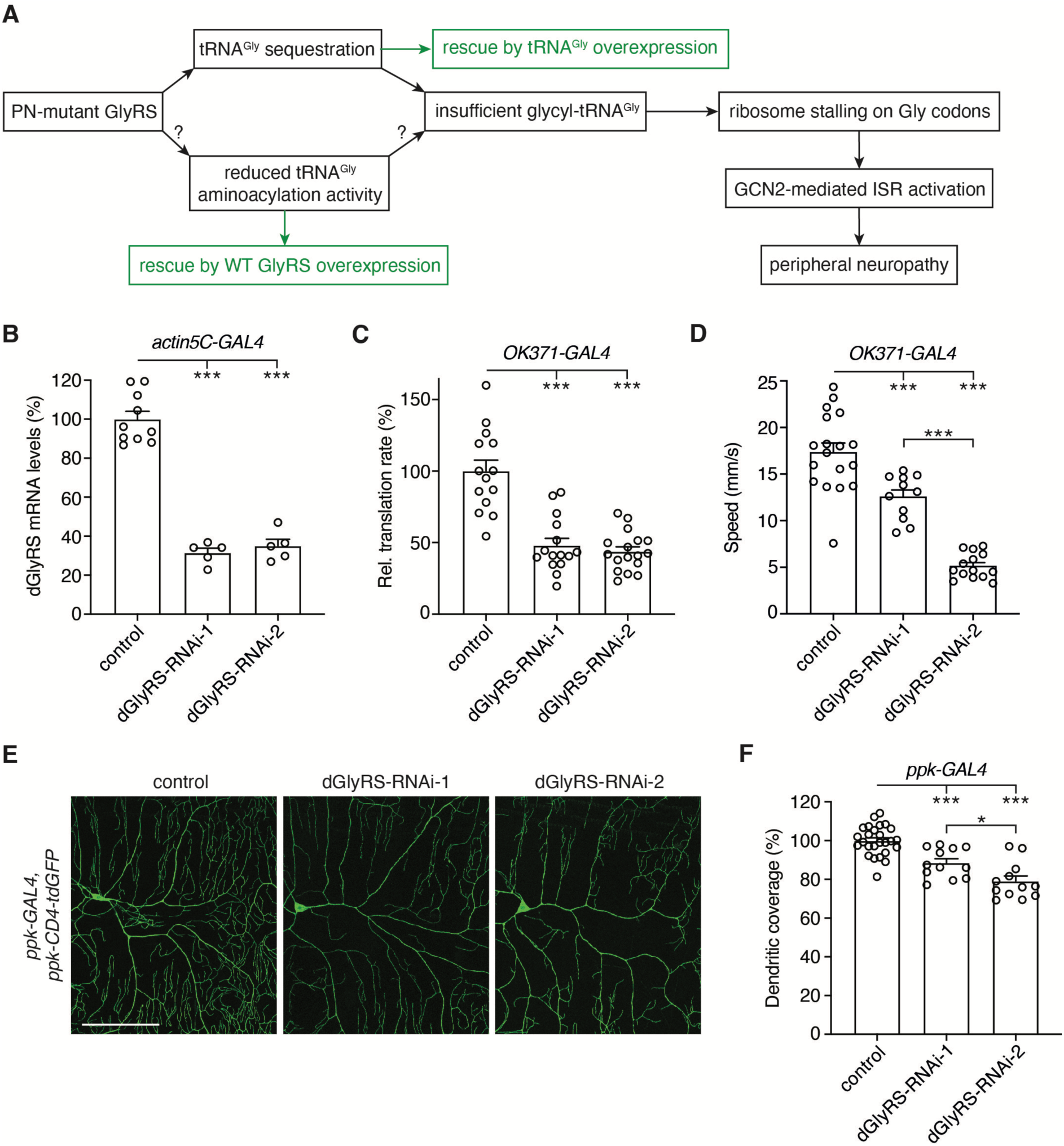
Transgenic knock-down of endogenous dGlyRS induces peripheral neuropathy phenotypes. A, Schematic representation of possible PN-GlyRS pathogenic mechanisms. Evidence for tRNA^Gly^ sequestration was previously reported for seven PN-GlyRS variants (12). Here, we evaluated whether some PN-GlyRS variants may – additionally or alternatively – cause disease through reducing tRNA^Gly^ aminoacylation activity. Optimal therapeutic approaches for each of the two mechanisms are shown in green. **B,** Endogenous dGlyRS mRNA levels as determined by quantitative real-time PCR (qPCR) on whole third instar larvae that ubiquitously (*actin5C-GAL4*) express one of two independent dGlyRS-RNAi transgenes. Data are shown as percentage (%) of driver-only control larvae. n = 5 to 10 samples per genotype, with each sample consisting of three larvae; ***p<0,0001 by one-way ANOVA with Tukey’s multiple comparisons test. **C,** Relative translation rate as determined by FUNCAT in motor neurons (*OK371-GAL4*) of third instar larvae expressing dGlyRS-RNAi transgenes. Data are shown as % of driver-only control larvae. n = 14 to 17 animals per genotype; ***p<0,0001 by Brown-Forsythe and Welch ANOVA with Dunnett’s T3 multiple comparisons test. **D,** Climbing speed in an automated negative geotaxis assay of 7-day-old female flies expressing dGlyRS-RNAi transgenes in motor neurons (*OK371-GAL4*). Data are shown as % of driver-only control flies. n = 11 to 18 groups of 10 flies per genotype; ***p<0,005 by Brown-Forsythe and Welch ANOVA with Dunnett’s T3 multiple comparisons test. **E,F,** Representative images (E) and quantification (F) of dendritic coverage (% of driver-only control) of class IV multidendritic sensory neurons in the larval body wall upon selective expression of dGlyRS-RNAi transgenes in these neurons (*ppk-GAL4*). n = 12 to 26 animals per genotype; *p<0.05; ***p<0.0005 by one-way ANOVA with Tukey’s multiple comparisons test. Scale bar: 100μm.

Human and mouse genetic evidence provide strong arguments against haplo-insufficiency as the mechanism underlying PN-GlyRS (1,16,17). However, because GlyRS functions as a homodimer, in heterozygous PN-GlyRS patients ∼25% of GlyRS dimers will be WT homodimers, ∼25% mutant homodimers, and ∼50% WT-mutant heterodimers, assuming an equal *GARS1* allelic expression and unaffected GlyRS dimerization. For PN-GlyRS mutations that result in a full loss of aminoacylation activity, this raises the possibility that a dominant negative mechanism may lead to an overall reduction of tRNA^Gly^ aminoacylation activity by 75%, because only WT homodimers would be able to aminoacylate tRNA^Gly^ (18).

Here, we evaluated the hypothesis that a subset of PN-GlyRS mutations may induce peripheral neuropathy through a dominant negative effect on tRNA^Gly^ aminoacylation. The plausibility of a dominant negative mechanism underlying some PN-GlyRS mutations is suggested by our finding that knock-down of endogenous *Drosophila* GlyRS (dGlyRS) in motor or sensory neurons is sufficient to induce peripheral neuropathy-related phenotypes. Furthermore, genetic experiments using three previously characterized PN-GlyRS *Drosophila* models (10) reveal that for two mutations that result in severely reduced or complete loss of aminoacylation activity (G240R and G526R), a dominant negative effect on aminoacylation activity might contribute to peripheral neuropathy, in addition to tRNA^Gly^ sequestration.

However, newly generated *Drosophila* PN-GlyRS models with *a priori* lower likelihood to sequester tRNA^Gly^ strongly argue for tRNA^Gly^ sequestration as a unifying mechanism underlying PN-GlyRS, and against a dominant negative mechanism. Indeed, expression of two PN-GlyRS variants that do not alter overall GlyRS protein charge (S211F and H418R) in *Drosophila* induced peripheral neuropathy phenotypes and inhibited global protein synthesis, highly reminiscent of our previously characterized PN-GlyRS models (10). Using *Drosophila* genetic experiments and biochemical assays, we show that tRNA^Gly^ sequestration by these newly characterized hGlyRS variants triggers peripheral neuropathy phenotypes. Finally, even high-level expression of the only reported PN-GlyRS variant that removes a positive charge from the GlyRS protein (K456Q) does not induce peripheral neuropathy phenotypes, and hGlyRS-K456Q display WT-like aminoacylation activity, suggesting that K456Q may not be a pathogenic mutation. Together, our results indicate that tRNA^Gly^ sequestration is a unifying pathogenic mechanism underlying PN-GlyRS. This implies that elevating tRNA^Gly^ levels may constitute a therapeutic approach for all PN-GlyRS patients, irrespective of their specific mutation.

## MATERIALS AND METHODS

### Drosophila genetics

Flies were housed in a temperature-controlled incubator with 12:12 h on/off light cycle at 25 °C. *OK371-GAL4* (Bl 26160), *ppk-GAL4* (Bl 32079), *actin5C-GAL4* (Bl 4414),and *Sca-GAL4* (Bl 6479) were obtained from the Bloomington *Drosophila* Stock Center (Bloomington, IN, USA). The UAS-dGlyRS line (19) was kindly provided by L. Luo. The UAS-hGlyRS-WT, UAS-hGlyRS-E71G, UAS-hGlyRS-G240R, and UAS-hGlyRS-G526R lines were previously described (10). For imaging of Class IV multidendritic sensory neurons, *ppk-GAL4* was recombined with *ppk-CD4-tdGFP* (20). dGlyRS-RNAi-1 corresponds to NIG 6778R-1, and dGlyRS-RNAi-2 corresponds to VDRC KK 106748. These transgenic RNAi lines were obtained from the National Institute of Genetics (NIG, Japan) and the Vienna Drosophila Resource Center (VDRC, Vienna, Austria), respectively. tRNA^Gly-GCC^ BAC and tRNA^Gly-GCC^ scramble transgenic lines were previously described (12). For ubiquitous expression of the UAS-hGlyRS transgenes from the adult stage onwards, the GAL80 target system was used (21). Fly crosses were cultured at 18 °C and 1-day-old adult flies carrying the *tub-GAL4*, tub-GAL80^ts^ and UAS-hGlyRS transgenes were shifted to 29 °C to induce transgene expression.

### Generation of hGlyRS transgenic lines

For generation of UAS-hGlyRS-S211F, UAS-hGlyRS-H418R, and UAS-hGlyRS-K456Q transgenic lines, full-length human GARS1 cDNA was purchased from Origene (Rockville, MD, USA) and subcloned into the pGEM^®^-T vector. Mutations were introduced via site-directed mutagenesis into the human GARS1 cDNA using the Q5^®^ Site-Directed Mutagenesis Kit (New England Biolabs, Ipswich, MA, USA). Sequences of primers used for cloning and mutagenesis of hGARS1 cDNA are listed in Supplementary Table 1. Next, hGARS1 cDNA was cloned into the pUAST-AttB transformation vector. Constructs were sequence verified and injected into *Drosophila* embryos by Genetivision (Houston, TX, USA) for site-specific PhiC31-mediated insertion in VK37 or VK31 landing sites for second or third chromosome insertions, respectively (22,23). Similar to the previously generated UAS-hGlyRS transgenes (10), the newly generated UAS-hGlyRS transgenes enable expression of either the cytosolic or mitochondrial hGlyRS isoform, depending on the translation start site used.

### Adult offspring frequency

For the determination of adult offspring frequency, the number of flies eclosing was counted for each genotype. A strong ubiquitous (*actin5C-GAL4*) driver was used to express the transgenes and flies were kept at a 12 h on/off light cycle at 25 °C. Eclosed flies were counted daily. At least 77 flies per genotype were counted.

### Scoring of scutellar bristle phenotypes

Bristle phenotypes were induced by expression of dGlyRS-RNAi-2 in sensory organ precursor cells using the *Sca-GAL4* driver, and we evaluated whether these bristle phenotypes could be rescued by simultaneous expression of either hGlyRS-WT or hGlyRS-K456Q. For quantification of bristle phenotypes, anterior and posterior scutellar bristles were examined and the number of normal, small or missing bristles was scored and the relative percentages were calculated.

### Analysis of protein synthesis by *in vivo* FUNCAT

For FUNCAT analysis of *de novo* protein synthesis (10,24,25), 24 h egg collections were performed and animals were raised on Jazz-Mix *Drosophila* medium (Fisher Scientific) at 25 °C. 72 h after egg laying, larvae were transferred to 4 mM azidonorleucine (ANL)-containing medium for 48 h (Iris Biotech, HAA1625). Larval central nervous system (CNS) was dissected in ice-cold HL3 solution and fixed in 4% paraformaldehyde (PFA) for 30 min at room temperature (RT). After fixation, the CNSs were washed 3 x 10 min with PBST (1x PBS pH 7.2 containing 0.2% Triton X-100) and 3 x 10 min with PBS pH 7.8. Metabolically labeled proteins were tagged by ‘click chemistry’ (Copper-Catalyzed (3+2)-Azide-Alkyne-Cycloaddition Chemistry (CuAAC)) using the fluorescent TAMRA tag. The FUNCAT reaction mix was assembled in a defined sequence of steps. Triazole ligand (200 µM, Sigma-Aldrich, 678937), TAMRA-alkyne tag (0.2 µM, Invitrogen, T10183), TCEP solution (400 µM, Sigma-Aldrich, C4706) and CuSO4 solution (200 µM, Sigma-Aldrich, C8027) were added to PBS pH 7.8. After each addition the solution was mixed thoroughly for 10 sec and at the end for 30 sec using a high-speed vortex. Larval CNSs were incubated with 500 μL of FUNCAT reaction mix overnight at 4 °C on a rotating platform. The next day, the CNSs were washed 3 times with PBS-Tween (1x PBS pH 7.2 containing 1% Tween-20) and 3 times with PBST for 10 min. Finally, larval CNSs were mounted in VectaShield mounting medium (Biozol, VEC-H-1000-CE) and stored at 4 °C until imaging using a Leica SP8 laser scanning confocal microscope. For image acquisition, identical confocal settings were used for all samples of a given experiment. Fluorescence intensities were quantified using ImageJ/FIJI software (National Institutes of Health). Mean intensity of all cells within one motor neuron cluster from each ventral nerve cord were used as single data points for statistical analysis.

### Analysis of protein synthesis by *in vivo* THRONCAT

For THRONCAT analysis of *de novo* protein synthesis (26), 24 h egg collections were performed and larvae were raised on Jazz-Mix *Drosophila* medium (Fisher Scientific) at 25 °C. 72 h after egg-laying, larvae were transferred to medium containing 4 mM ß-ethynylserine (ßES) for 48 h. Larval CNS was dissected in ice-cold HL3 solution and fixed in 4% PFA for 30 min at RT. After fixation, the CNSs were washed 3 x 10 min with PBST (1x PBS pH 7.2 containing 0.2% Triton X-100) and 3 x 15 min with PBS pH 7.2. CuAAC ‘click chemistry’ was performed using the fluorescent TAMRA tag and the reaction mix was assembled as follows: Tris-hydroxypropyltriazolylmethylamine (THPTA, Sigma-Aldrich, 762342) ligand (200 μM), TAMRA-azide tag (0.2 μM, Invitrogen, T10182), CuSO4 solution (4 mM), and sodium ascorbate solution (40 mM, Sigma-Aldrich, A7631) were added to PBS pH 7.2. After each addition the solution was mixed thoroughly for 10 sec and at the end for 30 sec using a high-speed vortex. CNSs were incubated with 500 µL of the reaction mix overnight at 4 °C on a rotating platform. The next day, CNSs were washed three times with PBS-Tween (1x PBS pH 7.2 containing 1% Tween-20) and three times with PBST for 15 min. Next, samples were blocked for 1 h in blocking buffer (10% goat serum in PBST) at RT before incubating with antibodies against hGlyRS (rabbit polyclonal; 1:200; Proteintech, 15831-AP) in blocking buffer at 4 °C overnight on a rotating platform. The next day, CNSs were washed three times with PBST for 15 min before incubation with Alexa Fluor™ 647-conjugated goat-anti-rabbit IgG antibodies (1:500; Thermo Fisher, A27040) in blocking buffer for 3 h at RT. After three washes with PBST for 15 min, CNSs were mounted in VectaShield mounting medium (Biozol, VEC-H-1000-CE) and stored at 4 °C until imaging using a Leica SP8 laser scanning confocal microscope. For image acquisition, identical confocal settings were used for all samples of a given experiment. Fluorescence intensities were quantified using ImageJ/FIJI software (National Institutes of Health). Mean intensity of all cells within one motor neuron cluster from each ventral nerve cord was determined, as well as the mean intensity of 5 cells in close proximity of the motor neuron cluster that do not express hGlyRS transgenes. The mean intensity of the motor neuron cluster was normalized to the mean intensity of the 5 surrounding cells and used as single data point for statistical analysis.

### Analysis of larval muscle innervation

To analyze synapse length on muscle 8, third instar larvae were dissected in HL3 buffer and fixed in Bouin’s for 3 min at RT. After permeabilization and blocking (10% goat serum in PBST), immunostaining was performed with anti-Discs large 1 (mouse monoclonal; 1:200; DSHB 4F3), to visualize the postsynaptic side of the neuromuscular synapse. Images were taken of muscle 8 in abdominal segment 5 using a Leica SP8 laser scanning confocal microscope with 20x Plan-Apochromat objective (0.8 NA). Confocal sections were taken every 2 μm covering the entire 3D volume of the NMJ. Using Fiji software, maximum intensity projections of all images in a z-stack were used to measure the synapse length, using the ‘segmented line’ tool to draw a line through the middle of every branch of the NMJ. When multiple NMJ branches were present, the sum of the length of individual branches was used as synapse length. The mean of the length of the left and right NMJ on muscle 8 was used as one data point for statistical analysis. For innervation status of muscle 24, the presence of the NMJ on muscle 24 in abdominal segment 6 on both sides of the larvae was assessed and counted as present (one or both sides) or absent (neither side).

### Automated negative geotaxis assay to analyze motor performance

To evaluate motor performance, newly eclosed male or virgin female flies were collected and kept in groups of ten flies per vial until measurements. Flies were not anaesthetized 24 h before measurements. Seven-day-old flies were flipped to test tubes and their motor performance was analyzed in a rapid iterative negative geotaxis assay (RING-assay) as previously described (10). Five or six iterative movies were recorded using a Nikon Z50 camera. A python-based script was used to analyze every movie by tracking the position of each fly in each frame and tracking all flies, creating tracks and allowing to obtain the climbing speed (mm/s) over the course of 5 seconds. The speed of all flies within one tube was averaged. The three movies with the highest number of moving flies detected were then averaged and used as one data point for the respective genotype. Movies with less than three moving flies were excluded from analysis.

### Analysis of dendritic morphology of class IV multidendritic sensory neurons

Imaging of class IV multidendritic sensory neuron morphology was performed on living third instar larvae. Microscopy slides were prepared with two coverslips glued on as spacers with conventional superglue. Larvae were thoroughly washed in tap water and single chilled larvae were immobilized under a coverslip, which itself was glued onto the spacing coverslips to produce equal spacing and pressure on the larvae. Only dorsal class IV da neurons of larval segment A1 were imaged using a Leica SP8 laser scanning confocal microscope. For quantification, maximum intensity projections were processed with Adobe Photoshop, rotated and digitally placed under a transparent grid containing a crosshair. The crosshair was used to fit the grid onto the cell body of the class IV da neuron in order to divide the dendritic tile into four quadrants. The posterolateral quadrant of the imaged da neuron was chosen for quantification of dendritic coverage, as least imaging artifacts and interference of denticle bands were encountered in this area. For quantification within the posterolateral quadrant, a sector of 200 μm to the lateral and 280 μm to the posterior was defined, equaling 56,000 μm^2^ divided into 560 squares of 8 x 8 pixels each. Presence versus absence of a piece of dendrite in the defined boxes was scored and calculated as measure of relative dendritic coverage.

### Western blot analysis

For western blot analysis, *tub-GAL4*, tub-GAL80^ts^ was used to express the UAS-hGlyRS transgenes from the adult stage onwards. Flies were raised at 18 °C and 1-to 3-day-old adult flies carrying the *tub-GAL4*, tub-GAL80^ts^ and UAS-hGlyRS transgenes were shifted to 29 °C to induce transgene expression. After five days of transgene expression, flies were snap frozen in liquid nitrogen and fly heads were collected. Protein extracts were prepared from 30 fly heads per sample by homogenization in extraction buffer (50 mM Tris/HCl pH 7.4, 150 mM KCl, 0.25 M sucrose, 5 mM MgCl2, and 0.5% Triton X-100 containing 1 U complete mini protease inhibitor cocktail (Roche)). Next, samples were centrifugated at 4 °C for 30 min at 13,000 x g to pellet cell debris. Samples were diluted to a concentration of 100 µg/mL. 20 µL per sample was separated on a 10% SDS-PAGE gel and transferred to a polyvinylidene difluoride (PVDF) membrane using the Trans-Blot Turbo Mini 0.2 µm PVDF Transfer Pack (Bio-Rad) for 10 min at 25 V, 2.5 A on a Trans-Blot^®^ Turbo Transfer System (Bio-Rad). The blotted membranes were blocked in 5% milk in TBS-T (20 mM Tris Base pH 7.6, 140 mM NaCl, 0.1% Tween-20) for 1 h at RT before overnight incubation at 4 °C with antibodies against hGlyRS (rabbit polyclonal; 1:1000; Proteintech, 15831-AP) or histone H3 (rabbit polyclonal; 1:10,000; Abcam, Ab1791). Membranes were subsequently incubated with anti-rabbit IgG coupled to horseradish peroxidase (goat polyclonal; 1:10,000; Invitrogen, 31460) for 1 h at RT to visualize proteins. Membranes were developed with enhanced chemiluminescence (ECL™ Prime Western Blotting System, Cytiva, RPN2232) and scanned on a Bio-Rad Universal Hood III Imager (Bio-Rad). Band intensities were quantified using ImageJ/FIJI (National Institutes of Health) by drawing a rectangle around the specific band, obtaining the peak pattern of the rectangle and measuring the area under the peak that represents the specific band.

### Isolation of total RNA and quantitative real-time PCR analysis

#### Quantification of dGlyRS transcript levels

dGlyRS-RNAi or dGlyRS transgenes were expressed with a ubiquitous driver (*actin5C-GAL4*). Three (dGlyRS-RNAi) or five (dGlyRS) third instar larvae were pooled per sample and total RNA was extracted using Trizol reagent (Life Technologies) and Direct-zol RNA Microprep kit (R2062, Zymol) according to the manufacturer’s instructions. Samples were DNase treated and 2500 ng RNA was subsequently used for reverse transcription to obtain cDNA, using the SensiFAST cDNA synthesis kit (BioLine). cDNA samples were then used as templates for real-time PCR assays performed on a Bio-Rad CFX system using the SensiFAST SYBR No-ROX Kit (BioLine). Beta-actin (dGlyRS-RNAi) or alpha-tubulin (dGlyRS) mRNA was used as reference gene for normalization. Experiments included no-reverse transcriptase and no-template controls for each primer pair. Primer sequences can be found in Supplementary Table 2. Data were analyzed using the ΔΔCt calculation method.

#### Quantification of hGlyRS transcript levels

hGlyRS transgenes were expressed with a ubiquitous driver from the adult stage onwards (*tub-GAL4*, tub-GAL80^ts^). Heads of 30 1-to 3-day-old adult flies were collected and snap frozen in liquid nitrogen. Heads were ground to a powder with a pestle and total RNA was extracted using Trizol reagent (Life Technologies) and Direct-zol RNA Microprep kit (R2062, Zymol) according to the manufacturer’s instructions. 500 ng RNA was used for cDNA synthesis using the SensiFAST cDNA synthesis kit (BioLine). cDNA samples were then used as templates for real-time PCR assays performed on a Bio-Rad CFX system using the SensiFAST SYBR No-ROX Kit (BioLine). Alpha-tubulin mRNA was used as a reference gene for normalization. Experiments included no-reverse transcriptase and no-template controls for each primer pair. Primer sequences can be found in Supplementary Table 2. Data were analyzed using the ΔΔCt calculation method.

### *In vitro* aminoacylation assay

*In vitro* transcription of *Drosophila* tRNA^Gly-GCC^ and aminoacylation with was performed as described (27). Briefly, tRNA^Gly-GCC^ and the respective recombinant hGlyRS variant (WT or K456Q) were mixed in 10:1 molar ratio (10 µM:1µM) and incubated for 15 min at 37 °C in 60 mM Tris-HCl pH 7.5, containing 10 mM KCl, 12.5 mM MgCl2, 1 mM spermidine, 1.5 mM DTT, 1.5 mM ATP, 1 mM glycine and 0.5 µL SUPERase·In™ RNase Inhibitor (Invitrogen) in a total reaction volume of 20 µL. The reaction was terminated by dilution up to 300µl with 0.3 M NaOAc pH 4.5, containing 10 mM EDTA. The tRNA was precipitated with 1 volume isopropanol and resuspended in 10 mM NaOAc pH 4.5 (note that the ester bond is stable in acidic pH).

To distinguish between aminoacylated and uncharged tRNA species, we used the splinted ligation reaction that generates amino acid-bridged chimeric RNA molecules to a 5’-phosphorimidazolide-activated oligoribonucleotide (27), thereby following the reaction scheme and using the same RNA:DNA oligonucleotides used to extend the 5’ and 3’ ends of the tRNA as described (28). The oligonucleotide with 5’-phosphorimidazolide ligates preferably aminoacyl-tRNA entities with more than 3 orders of magnitude higher efficacy than the non-amonoacylated tRNAs. tRNA^Gly-GCC^ recovered after the aminoacylation reaction was incubated with the 5’-phosphorimidazolide oligonucleotide for 30 min at 25 °C in 100 mM MES buffer pH 5.5, containing 2.5 mM MgCl_2_, in the presence of 50 mM 1-(2-hydroxyethyl)imidazole (HEI) pH 6.5 catalyst, followed by an extended incubation at 4 °C overnight. The different tRNA species were separated on a 10% denaturing acidic PAGE (8M urea and 0.1 M NaOAc pH 5) and visualized by SYBR^TM^ Gold Nucleic Acid Stain and quantified using ImageJ software. The percentage of the aminoacyl-tRNA was determined as the ratio to the total (aminoacyl-tRNA and non-aminoacyted tRNA) and data are represented in Figure 3K as aminoacylation activity relative to hGlyRS-WT (100%).

**Figure 2:**
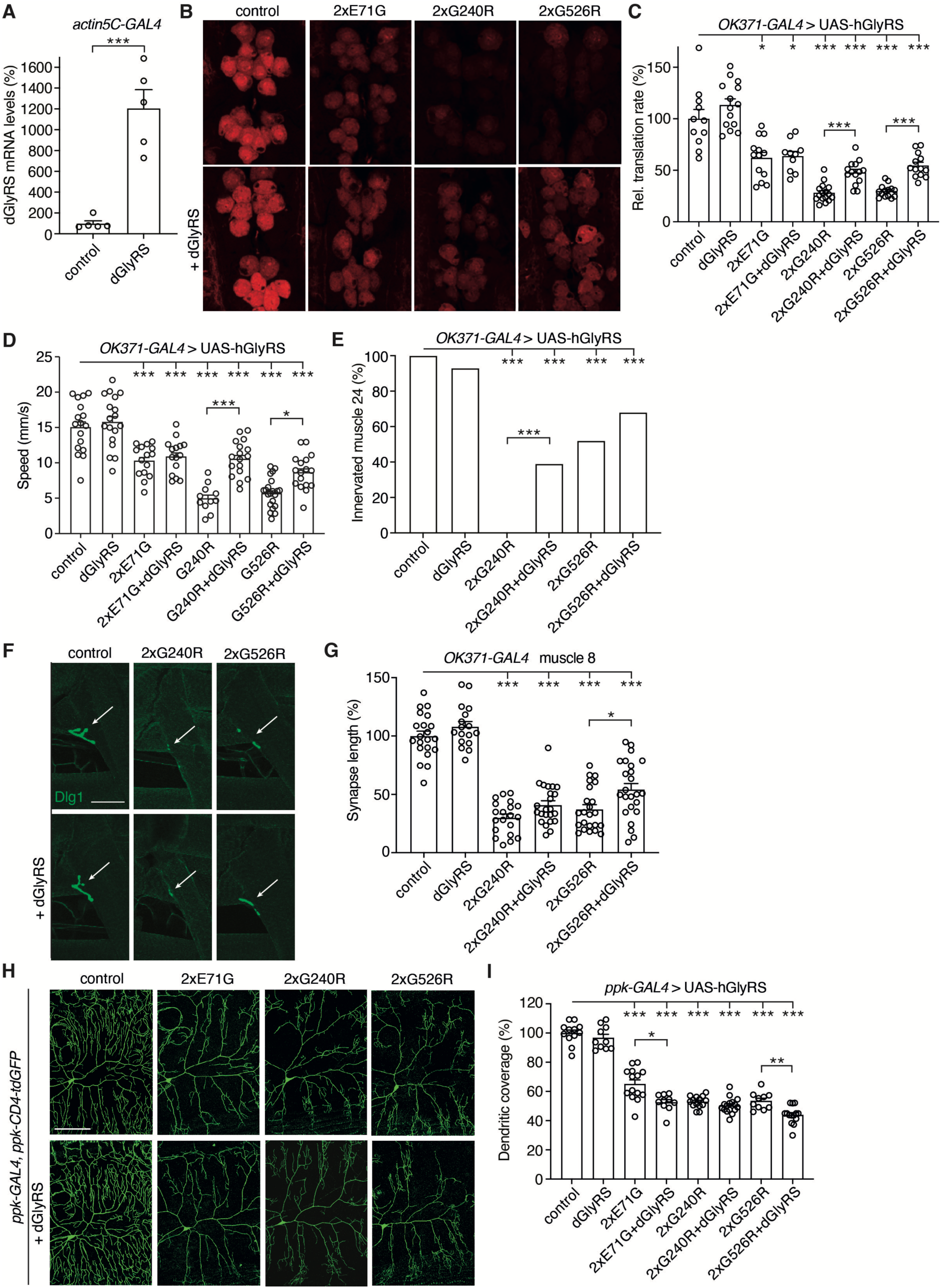
Effect of dGlyRS overexpression on protein synthesis and peripheral neuropathy phenotypes of PN-GlyRS *Drosophila* models. A, dGlyRS mRNA levels as determined by qPCR on whole third instar larvae that ubiquitously (*actin5C-GAL4*) express a dGlyRS transgene. Data are shown as percentage (%) of driver-only control larvae. n = 5 samples per genotype, with each sample consisting of five larvae; ***p<0.005 by unpaired t-test with Welch’s correction. **B,C,** Representative images (B) and quantification (C) of relative translation rate (% of driver-only control) as determined by FUNCAT in motor neurons (*OK371-GAL4*) of larvae expressing E71G, G240R, or G526R hGlyRS (2x: two transgene copies), in the presence or absence of *Drosophila* GlyRS (dGlyRS) co-overexpression. n = 10 to 16 animals per genotype; *p<0.05; ***p<0.005 by Brown-Forsythe and Welch ANOVA with Dunnett’s T3 multiple comparisons test. Scale bar: 10μm. **D,** Climbing speed in a negative geotaxis assay of 7-day-old female flies expressing hGlyRS transgenes in motor neurons (*OK371-GAL4*), in the presence or absence of dGlyRS co-overexpression. n = 11 to 21 groups of 10 flies per genotype; *p<0.05; ***p<0.0005 by one-way ANOVA with Sidak’s multiple comparisons test. **E,** Percentage of larvae with innervated muscle 24. hGlyRS transgenes were expressed in motor neurons (*OK371-GAL4*), in the presence or absence of dGlyRS co-overexpression. n = 14 to 26 animals per genotype; ***p<0.005 by Fisher’s exact test. **F,G,** Representative images (F) and quantification (G) of neuromuscular synapse length on distal muscle 8 of larvae expressing hGlyRS transgenes in motor neurons (*OK371-GAL4*), in the presence or absence of dGlyRS co-overexpression. n = 17 to 24 animals per genotype; *p<0.05; ***p<0.0001 by one-way ANOVA with Sidak’s multiple comparisons test. Scale bar: 50μm. **H,I,** Representative images (H) and quantification (I) of dendritic coverage (% of driver-only control) of class IV multidendritic sensory neurons in the larval body wall upon selective expression of hGlyRS transgenes in these neurons (*ppk-GAL4*), in the presence or absence of dGlyRS co-overexpression. n = 10 to 16 animals per genotype; *p<0.05; **p<0.01; ***p<0.0001 by Brown-Forsythe and Welch ANOVA with Dunnett’s T3 multiple comparisons test. Scale bar: 100μm.

**Figure 3:**
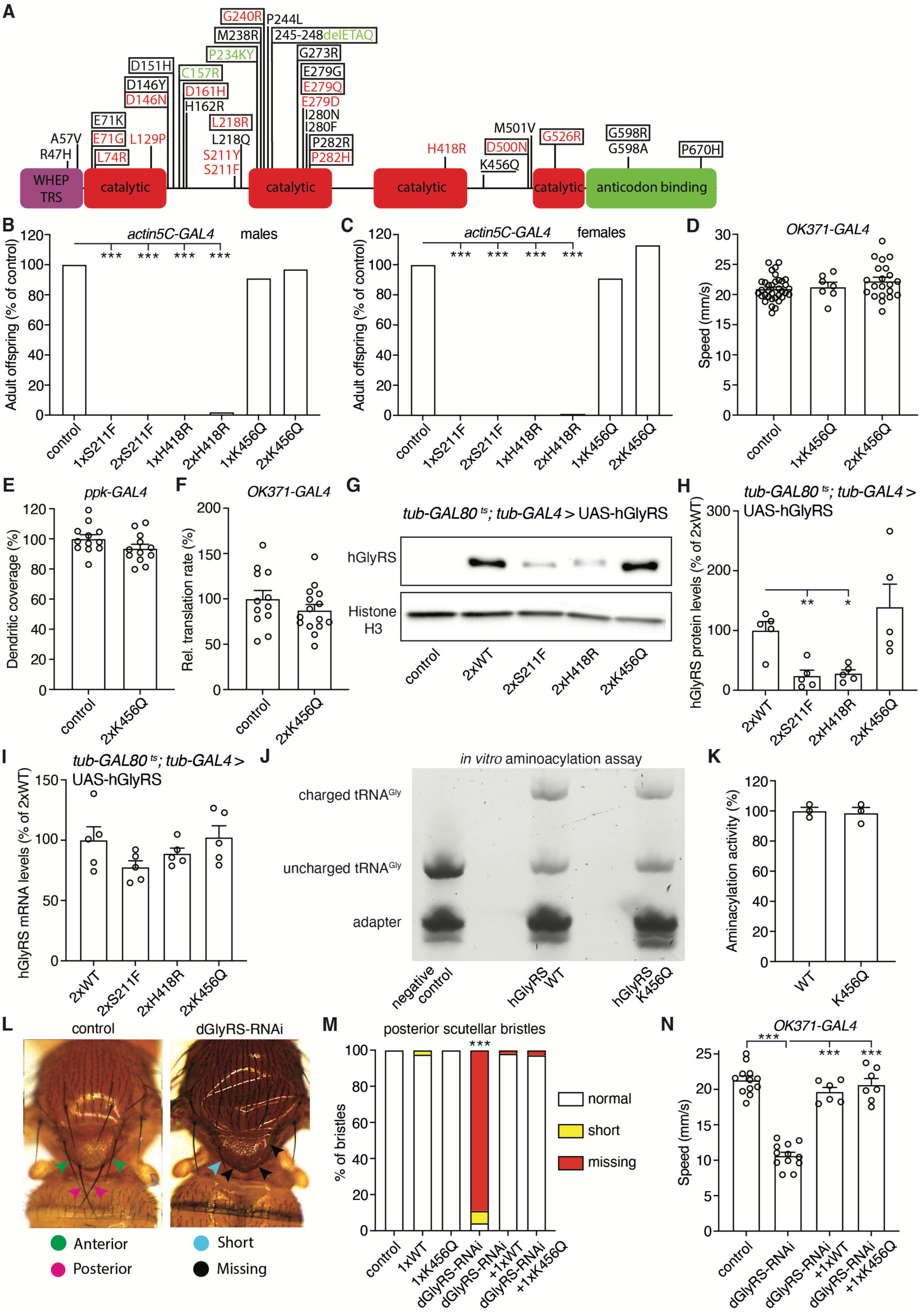
hGlyRS-K456Q does not induce peripheral neuropathy phenotypes in *Drosophila* and retains aminoacylation activity. A, Schematic representation of PN-GlyRS mutations. Mutations that segregate with disease in a family (strongest genetic evidence for disease causality) are labeled in red, mutations found in single patients are in black, and mutations in PN-aaRS mouse models are in green. Mutations that increase the charge of GlyRS (add positive charge) are boxed, while the single mutation that reduces the charge is underlined. The WHEP-TRS, catalytic, and anticodon binding domains are indicated. The positions of the mutations refer to the cytosolic isoform. Figure adapted from (1). **B,C,** Adult offspring frequency (% of driver-only control) of male (B) and female (C) flies ubiquitously (*actin5C-GAL4*) expressing one or two copies of hGlyRS S211F, H418R, or K456Q transgenes. n = 77 to 494 (B) and 144 to 523 (C) animals per genotype; ***p<0.0001 by Fisher’s exact test. **D,** Negative geotaxis climbing speed of 7-day-old female flies expressing one or two copies of hGlyRS-K456Q transgene in motor neurons (*OK371-GAL4*), as compared to driver-only control. n = 7 to 35 groups of 10 flies per genotype; p = not significant by one-way ANOVA with Tukey’s multiple comparisons test. **E,** Quantification of dendritic coverage (% of driver-only control) of class IV multidendritic sensory neurons in the larval body wall upon selective expression of hGlyRS-K456Q in these neurons (*ppk-GAL4*). n = 12 animals per genotype; p = 0.11 by unpaired t-test. **F,** Relative translation rate (% of driver-only control) as determined by FUNCAT in motor neurons (*OK371-GAL4*) of larvae expressing hGlyRS-K456Q. n = 12 to 15 animals per genotype; p = 0.26 by unpaired t-test. **G,H,** Representative image (G) and quantification (H) of western blot to evaluate hGlyRS protein levels in adult heads of flies ubiquitously expressing WT, S211F, H418R, or K456Q hGlyRS from the adult stage onwards (*tub-GAL80^ts^; tub-GAL4*), harvested five days after induction of transgene expression. Control is driver-only. Histone H3 was used as loading control. Data are shown as % of 2xWT. n = 5 independent samples per genotype; *p<0.05; **p<0.01 by Brown-Forsythe and Welch ANOVA with Dunnett’s multiple comparisons test. **I,** hGlyRS mRNA levels as determined by quantitative real-time PCR on heads of flies ubiquitously expressing WT, S211F, H418R, or K456Q hGlyRS from the adult stage onwards (*tub-GAL80^ts^; tub-GAL4*), harvested five days after induction of transgene expression. Data are shown as percentage (%) of flies expessing two copies of hGlyRS-WT. n = 5 independent samples per genotype; p = not significant by one-way ANOVA with Sidak’s multiple comparisons test. **J,K,** Representative image (J) and quantification (K) of an *in vitro* aminoacylation assay in which tRNA^Gly-GCC^ is incubated with glycine and recombinant hGlyRS-WT or hGlyRS-K456Q for 15 minutes, followed by adapter ligation selectively to glycyl-tRNA^Gly^ (and not to uncharged tRNA^Gly^) and separation by denaturing acidic PAGE. The ratio of glycyl-tRNA^Gly^ to total tRNA^Gly^ was determined and represented in (J) as tRNA^Gly^ aminoacylation activity relative to hGlyRS-WT (100%). n = 3 per hGlyRS variant; p = 0.77 by unpaired t-test. **L,** Representative images of bristle phenotypes induced by dGlyRS knock-down in sensory organ precursor cells (*Sca-GAL4*). Anterior and posterior scutellar bristles (indicated by green and pink arrowheads, respectively) are always long in controls, but often short (blue arrowhead) or missing (black arrowheads) in *Sca-GAL4*>UAS-dGlyRS-RNAi flies. **M,** Quantification of the percentage of normal (long), short, or missing posterior scutellar bristles in the presence or absence of dGlyRS knock-down, and with or without simultaneous co-expression of hGlyRS-WT or hGlyRS-K456Q. n = 36 to 118 bristles per genotype; ***p<0.0001 by Fisher’s exact test. **N,** Negative geotaxis climbing speed of 7-day-old female flies expressing dGlyRS-RNAi in motor neurons (*OK371-GAL4*), with or without simultaneous co-expression of hGlyRS-WT or hGlyRS-K456Q and compared to driver-only control. n = 6 to 12 groups of 10 flies per genotype; ***p<0.0001 by one-way ANOVA with Sidak’s multiple comparisons test. Note that in figure panels 3E and 4C, the data points of the driver-only control are the same, because they were generated in the same larger experiment of which the results are dispalyed in different panels for the sake of clarity of presentation.

### Expression and purification of recombinant hGlyRS proteins

Two PN-GlyRS variants (S211F and H418R) were expressed with two tags (6xHis and SUMO tag) in *E. coli* Rosetta strain and purified as described previously for other PN-GlyRS variants (12). Briefly, crude extracts were incubated with *HisPur Ni-NTA resin* (Thermo) for 30 minutes at 4 °C, followed by multiple washing steps with 20 mM HEPES-NaOH pH 7.4, containing 500 mM NaCl and 10 mM imidazole, and two washing steps with imidazole concentration increased to 20 mM. The PN-GlyRS variants were dissolved from the Ni-NTA resin by cleavage after the SUMO tag with 0.5 mg/mL of ULP overnight at 4 °C. The released PN-GlyRS variants in the supernatant were concentrated and the dimer and monomer forms were fractionated using size-exclusion chromatography on *Superdex 200*.

### Determination of *K*_on_ and *K*_off_ values

The *K*_on_ values for the cognate tRNA^Gly-GCC^ were determined by monitoring the quenching of intrinsic PN-GlyRS tryptophan residues as described previously (12). tRNA^Gly-GCC^ was generated via in-vitro T7 transcription and purified via 10% polyacrylamide gel electrophoresis. Briefly, 750 nM of each PN-GlyRS variant was incubated with different tRNA concentrations, ranging from 0-1µM, in 25 mM sodium acetate buffer pH 6.0 containing 10 mM MgCl_2_, 5 mM DTT, in a final volume of 50µL. The Trp emission was recorded at 350 nm (excitation 280 nm) at 37 °C on a *TECAN Spark* plate-reader. Binding curves were recorded in multiple replicates and fitted to exponential decay functions and quantified with *OriginPro*.

For determination of the *K*_off_ values, to each of the above reactions containing different (0.1-1µM) tRNA^Gly^ concentrations, 1 mM glycine and 1 mM ATP were added. Fluorescence spectra (emission at 350 nm and excitation at 280 nm) were recorded over 5 min, step 1 s. Spectra were fitted in *OriginPro* to an exponential function.

### Statistics

All data are presented as mean ± standard error of the mean (SEM) and differences were considered significant when p<0.05. Animals or samples were assigned to the various experimental groups based on their genotype. For a given experiment, samples from the different experimental groups were processed in parallel and analyzed in random order.

Before analysis, a Robust regression and Outlier removal method (ROUT) was performed to detect statistical outliers. This nonlinear regression method fits a curve that is not influenced by outliers. Residuals of this fit were then analyzed by a test adapted from the False Discovery Rate approach, to identify any outliers. All data points that were considered outliers were excluded from further data analysis. Normality and homoscedasticity of all data sets was assessed by a Shapiro-Wilk and Brown-Forsythe (F-test for t-tests) test, respectively. Subsequent statistical tests were only performed if all assumptions were met.

For comparison of normally distributed data of more than two groups, one-way ANOVA followed by Sidak’s or Tukey’s multiple comparisons *post hoc* test was used, provided that the SDs were not significantly different between groups (equal variance, evaluated by Brown-Forsythe test and Bartlett’s test). For comparison of normally distributed data of more than two groups with unequal variance, Brown-Forsythe and Welch ANOVA followed by Dunnett’s T3 multiple comparisons *post hoc* test was used. For comparison of not normally distributed data of more than two groups, Kruskal-Wallis test with Dunn’s multiple comparisons test was used. For comparison of normally distributed data from two groups with equal variance, unpaired t-test was used. Fisher’s exact test was used to analyze offspring frequency data, innervation status of larval muscle 24, and scutellar bristle phenotpes.

## RESULTS

### Endogenous dGlyRS knock-down inhibits protein synthesis and induces peripheral neuropathy phenotypes

To understand whether a loss-of-function scenario is a plausible mechanism for PN-GlyRS, we first evaluated whether a substantial reduction of dGlyRS levels, which consequently reduces tRNA^Gly^ aminoacylation, is sufficient to inhibit protein synthesis and induce peripheral neuropathy-related phenotypes. To do so, we used two independent transgenic RNAi lines that allow for knock-down of dGlyRS. To evaluate the efficiency of the dGlyRS-RNAi lines, we expressed the dGlyRS-RNAi transgenes ubiquitously (*actin5C-GAL4*), and quantified dGlyRS transcript levels in third instar larvae by quantitative real-time PCR. The two dGlyRS-RNAi lines reduced dGlyRS transcript levels by ∼69% and ∼65%, respectively (Figure 1B). Due to the lack of a dGlyRS antibody, it was not possible to determine the corresponding endogenous dGlyRS protein levels. Nevertheless, the reduction in dGlyRS transcript levels may be representative for a dominant negative scenario, in which the aminoacylation-active dGlyRS pool is reduced by up to 75%.

We next evaluated the effect of dGlyRS knock-down in motor neurons (*OK371-GAL4*), because motor neurons are the affected cell type in all *GARS1*-associated peripheral neuropathies. *In vivo* cell-type-specific FUNCAT (24,25) revealed a marked inhibition of nascent protein synthesis in motor neurons by ∼52% and ∼56% for the two independent dGlyRS-RNAi lines (Figure 1C). These data suggest that dGlyRS knock-down substantially inhibits tRNA^Gly^ aminoacylation, leading to depletion of cellular glycyl-tRNA^Gly^ levels and consequently inhibiting protein synthesis. To determine whether dGlyRS knock-down in motor neurons can trigger peripheral neuropathy-like phenotypes, we evaluated motor performance of 7-day-old adult flies using an automated negative geotaxis assay (10). This assay revealed that both dGlyRS-RNAi lines induced a significant reduction in climbing speed (Figure 1D). Finally, we evaluated the effect of dGlyRS knock-down on sensory neuron morphology, given that sensory neurons are the other affected cell type in *GARS1*-associated CMT (CMT2D). Selective knock-down of dGlyRS in class IV multidendritic sensory neurons in the body wall of third instar larvae (*ppk-GAL4*) significantly reduced their dendritic coverage for both dGlyRS-RNAi lines (Figure 1E,F). Together, these data indicate that a substantial reduction of dGlyRS levels is sufficient to inhibit global protein synthesis and induce peripheral neuropathy-related phenotypes.

### Effect of dGlyRS overexpression on protein synthesis and peripheral neuropathy phenotypes of PN-GlyRS *Drosophila* models

We previously generated and characterized PN-GlyRS *Drosophila* models that allow transgenic expression of three human PN-GlyRS mutant proteins: E71G, G240R, G526R (10). While we previously showed that each of these three hGlyRS variants sequester tRNA^Gly^ (12), we sought to explore whether a dominant negative effect may constitute an additional pathogenic mechanism. This is particularly relevant for the G240R and G526R mutations, which severely reduce tRNA^Gly^ aminoacylation activity (10,29–31). In contrast, hGlyRS-E71G retains aminoacylation activity (10,29,30)(Supplementary Table 3). In case of a dominant negative scenario, increasing WT dGlyRS protein levels should increase the fraction and number of WT dGlyRS dimers and therefore would be expected to rescue the PN-GlyRS phenotypes. We therefore evaluated the effect of transgenic dGlyRS overexpression on inhibition of protein synthesis and peripheral neuropathy phenotypes in our PN-GlyRS *Drosophila* models. Ubiquitous (*actin5C-GAL*) expression of the dGlyRS transgene in third instar larvae resulted in a 12-fold increase in dGlyRS mRNA levels (Figure 2A), confirming robust transgene expression. As expected, *in vivo* cell-type specific FUNCAT revealed inhibition of protein synthesis induced by expression of hGlyRS E71G, G240R or G526R variants in third instar larval motor neurons (Figure 2B,C). Simultaneous overexpression of dGlyRS did not modify the protein synthesis defect induced by hGlyRS-E71G, but partially rescued the inhibition of translation induced by hGlyRS G240R and G526R (Figure 2B,C). Similarly, the adult motor performance defect induced by hGlyRS-E71G expression in motor neurons was not altered by simultaneous dGlyRS overexpression, while adult motor deficits induced by hGlyRS-G240R and hGlyRS-G526R expression were partially rescued by dGlyRS co-overexpression (Figure 2D).

Next, we investigated third instar larval neuromuscular junction (NMJ) phenotypes induced by expression of hGlyRS-G240R and hGlyRS-G526R in motor neurons (10). Expression of hGlyRS-G240R or hGlyRS-G526R in motor neurons frequently resulted in denervation of larval muscle 24, and this denervation phenotype was mitigated by simultaneous dGlyRS overexpression. This effect reached statistical significance for hGlyRS-G240R but not for hGlyRS-G526R (Figure 2E). Furthermore, the neuromuscular synapse length on larval muscle 8 was significantly reduced by expression of hGlyRS-G240R and hGlyRS-G526R in motor neurons. This defect was slightly but significantly improved by simultaneous dGlyRS overexpression for the hGlyRS-G526R variant but not for hGlyRS-G240R (Figure 2F,G). Finally, selective expression of each of the three hGlyRS mutants in class IV multidendritic sensory neurons significantly reduced their dendritic coverage. Simultaneous dGlyRS overexpression did not rescue this sensory neuron morphology defect. Rather, for the E71G and G526R mutants, dGlyRS co-overexpression slightly but significantly enhanced the sensory neuron morphology defect (Figure 2H,I). Together, these data indicate that the hGlyRS-E71G variant does not act through a dominant negative mechanism. In contrast, at least in motor neurons, the protein synthesis defect and peripheral neuropathy phenotypes induced by the G240R and G526R variants may in part be attributable to a dominant negative mechanism, in addition to tRNA^Gly^ sequestration.

### Generation and characterization of novel *Drosophila* PN-GlyRS models

Of the 35 human PN-GlyRS mutations reported to date, 20 (57%) add net positive charge to the GlyRS protein (by replacing a negatively charged amino acid by a neutral or positively charged amino acid, a neutral residue by a positively charged one or by deleting a negatively charged residue), 14 do not change the GlyRS protein charge, and only one mutation increases the net negative charge by replacing a positively charged residue by an uncharged residue (K456Q) (Figure 3A). Coincidentally, each of the three GlyRS mutations that we previously modelled in *Drosophila* increase the net positive charge to the protein (E71G, G240R, G526R). Although we previously showed that addition of positive charge to the GlyRS protein is not required for a mutation to induce tRNA^Gly^ sequestration *in vitro* (12), increasing the net positive charge is likely to promote tRNA^Gly^ sequestration. We therefore reasoned that if some PN-GlyRS mutations would act through a dominant negative mechanism, mutations that do not alter the net GlyRS charge or even increase the net negative charge are the most likely candidates. Hence, we generated three novel PN-GlyRS *Drosophila* models. We selected the S211F and H418R mutations, because (1) they do not change the net GlyRS charge, (2) they clearly segregate with disease in families (32,34) and (3) they result in loss of aminoacylation activity (29,33,35), which is a prerequisite for a dominant negative mechanism (Supplementary Table 3). In addition, we selected the K456Q mutation (35,37) because this is the only mutation reported that renders the overall GlyRS charge more negative (Figure 3A, Supplementary Table 3). After introduction of the respective mutations, we used site-specific transgenesis to integrate the hGlyRS transgenes in landing sites on the second and third chromosomes of *Drosophila*. To maximize the comparability between PN-GlyRS models, the same genomic landing sites were used as for our initial PN-GlyRS *Drosophila* models. Similar to the previous PN-GlyRS *Drosophila* models (10), the newly generated UAS-hGlyRS transgenes allow for expression of the cytosolic or mitochondrial hGlyRS isoform, depending on the translation start site used.

### hGlyRS-K456Q does not induce peripheral neuropathy phenotypes in *Drosophila* and retains aminoacylation activity

As a first step, we evaluated whether expression of the new hGlyRS variants in an otherwise WT *dGlyRS* background would induce peripheral neuropathy phenotypes and inhibit protein synthesis. While the expression of hGlyRS-S211F and hGlyRS- H418R clearly induced peripheral neuropathy phenotypes and inhibited protein synthesis (see below and Figure 4), this was not the case for hGlyRS-K456Q. Whereas ubiquitous expression (*actin5C-GAL4*) of hGlyRS-S211F and hGlyRS- H418R transgenes induced developmental lethality, ubiquitous expression of either one or two copies of hGlyRS-K456Q transgene did not reduce adult offspring frequency (Figure 3B,C). Furthermore, expression of one or two copies of hGlyRS- K456Q transgenes in motor neurons did not induce adult motor performance deficits (Figure 3D). Moreover, even high-level expression of hGlyRS-K456Q in class IV multidendritic sensory neurons did not induce dendritic morphology defects (Figure 3E). Finally, high-level expression of hGlyRS-K456Q did not affect *de novo* protein synthesis in larval motor neurons (Figure 3F). To ensure that the lack of peripheral neuropathy phenotypes following hGlyRS-K456Q expression is not attributable to reduced hGlyRS-K456Q protein levels, we performed western blotting using heads of flies expressing hGlyRS transgenes ubiquitously from the adult stage onwards, to circumvent developmental lethality (*tubulin-GAL80^ts^; tubulin-GAL4,* see Materials and Methods for details). This experiment revealed that hGlyRS-K456Q levels were comparable to hGlyRS-WT levels, whereas hGlyRS-S211F and hGlyRS-H418R protein levels were significantly reduced (Figure 3G,H). The reduced hGlyRS-S211F and hGlyRS-H418R protein levels are likely due to inhibition of global protein synthesis induced by these PN-GlyRS mutants (see below and Figure 4D), because the presence PN-GlyRS mutations did not affect hGlyRS transcript levels (Figure 3I). Collectively, our data indicate that even high-level expression of hGlyRS-K456Q did neither affect protein synthesis, nor induced peripheral neuropathy in *Drosophila*, indicating that the K456Q variant does not act through a gain-of-toxic function mechanism.

**Figure 4:**
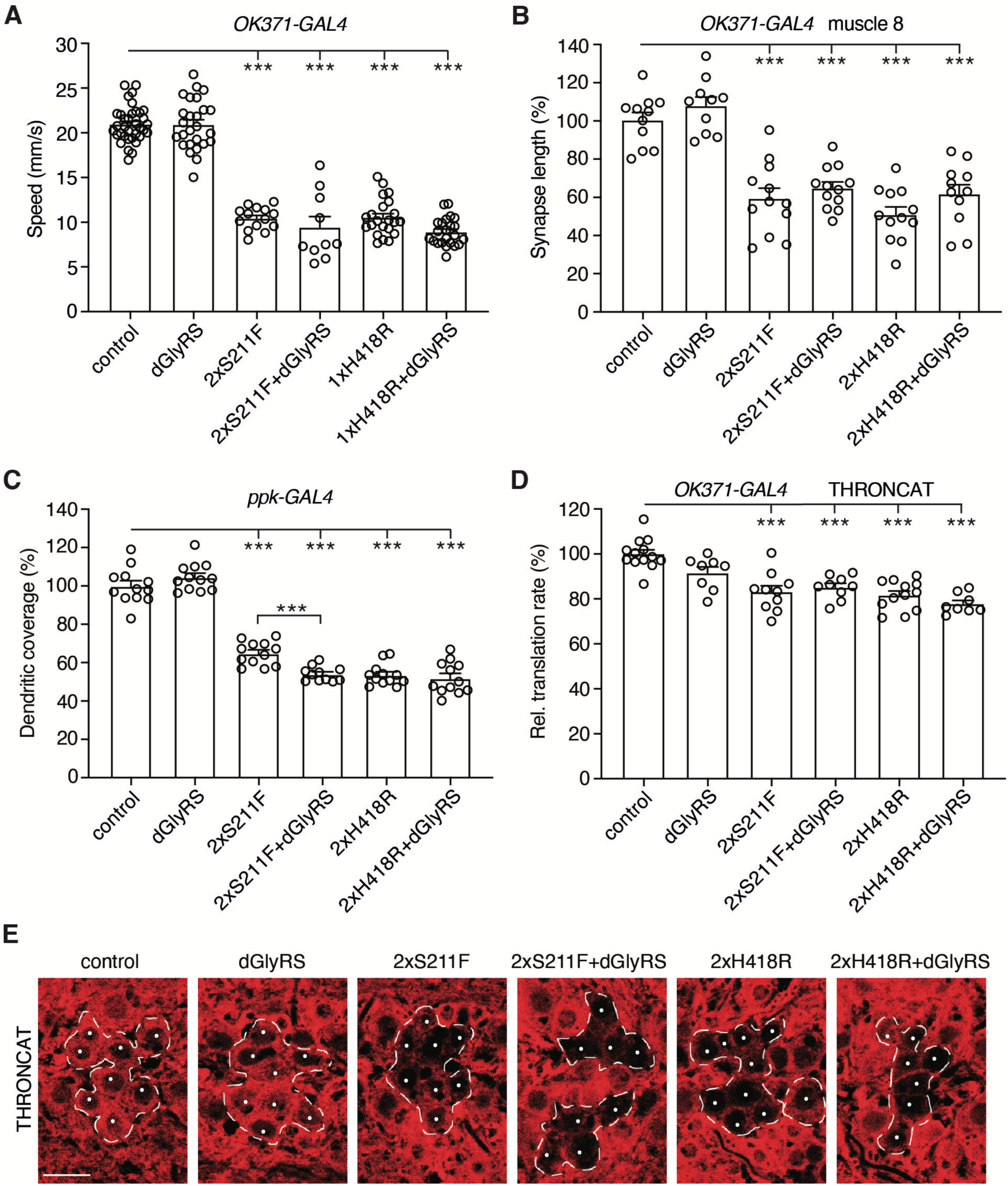
Peripheral neuropathy phenotypes induced by hGlyRS-S211F and hGlyRS- H418R are not attributable to a dominant negative mechanism. A, Negative geotaxis climbing speed of 7-day-old female flies expressing hGlyRS-S211F or hGlyRS-H418R transgenes in motor neurons (*OK371-GAL4*), in the presence or absence of dGlyRS co-overexpression. n = 10 to 35 groups of 10 flies per genotype; ***p<0.0001 by Kruskal-Wallis test with Dunn’s multiple comparisons test. **B,** Synapse length on distal muscle 8 of larvae expressing hGlyRS-S211F or hGlyRS-H418R transgenes in motor neurons (*OK371-GAL4*), in the presence or absence of dGlyRS co-overexpression. N = 10 to 12 animals per genotype; ***p<0.0001 by one-way ANOVA with Sidak’s multiple comparisons test. **C,** Quantification of dendritic coverage (% of driver-only control) of class IV multidendritic sensory neurons in the larval body wall upon selective expression of hGlyRS-S211F or hGlyRS-H418R transgenes in these neurons (*ppk-GAL4*), in the presence or absence of dGlyRS co-overexpression. n = 11 to 12 animals per genotype; ***p<0.005 by one-way ANOVA with Sidak’s multiple comparisons test. **D,E,** Quantification (D) and representative images (E) of relative translation rate (% of driver-only control) as determined by THRONCAT in motor neurons (*OK371-GAL4*) of larvae expressing hGlyRS-S211F or hGlyRS-H418R, in the presence or absence of dGlyRS co-overexpression. Motor neurons targeted by *OK371-GAL4* are delineated by a dashed line and nuclei of motor neurons are labeled by a white dot (E). n = 8 to 13 animals per genotype; ***p<0.0001 by one-way ANOVA with Sidak’s multiple comparisons test. Scale bar: 10μm.

Because the hGlyRS transgenes were expressed in a WT *dGlyRS* background, we wanted to exclude that hGlyRS-K456Q causes peripheral neuropathy in human CMT patients through a loss-of-function mechanism. We therefore evaluated the effect of the K456Q mutation on tRNA^Gly^ aminoacylation activity, using both an *in vitro* aminoacylation assay and *in vivo* genetic complementation experiments. The *in vitro* aminoacylation assay with recombinant hGlyRS proteins revealed that hGlyRS- K456Q displays tRNA^Gly^ aminoacylation activity similar to that of hGlyRS-WT (Figure 3J,K). Knock-down of dGlyRS in sensory organ precursor cells (*Sca-GAL4*>UAS- dGlyRS-RNAi-2) resulted in shorter or missing scutellar bristles, which are long in control animals (Figure 3L). Simultaneous expression of hGlyRS-WT fully rescued these bristle phenotypes (Figure 3M, Supplementary Figure 1), indicating the dGlyRS and hGlyRS are functional homologs. Consistent with the WT-like aminoacylation activity, expression of hGlyRS-K456Q also fully rescued dGlyRS-RNAi induced bristle phenotypes (Figure 3M, Supplementary Figure 1). Finally, the climbing defect induced by dGlyRS knock-down in motor neurons (*OK371-GAL4*) was also fully rescued by simultaneous expression of either hGlyRS-WT or hGlyRS-K456Q (Figure 3N). Together, our data show that hGlyRS-K456Q exhibits WT-like tRNA^Gly^ aminoacylation activity and indicate that the K456Q mutation may not be pathogenic for PN-GlyRS (see discussion).

### Peripheral neuropathy phenotypes induced by hGlyRS-S211F and hGlyRS- H418R are not attributable to a dominant negative mechanism

We next asked whether a dominant negative mechanism may underlie peripheral neuropathy phenotypes induced by hGlyRS-S211F and hGlyRS-H418R expression. Therefore, we evaluated the effect of dGlyRS co-overexpression, because increasing dGlyRS levels is expected to induce a phenotypic rescue in case of a dominant-negative mechanism. Expression of hGlyRS-S211F or hGlyRS-H418R in motor neurons reduced the climbing speed of adult flies in the negative geotaxis assay by ∼50% (Figure 4A). Strikingly, simultaneous dGlyRS overexpression did not rescue this motor performance defect. Expression of hGlyRS-S211F and hGlyRS-H418R in motor neurons also reduced the neuromuscular synapse length on larval muscle 8 by ∼41% and ∼49%, respectively (Figure 4B). Again, simultaneous dGlyRS overexpression did not significantly rescue this NMJ morphology defect. Selective expression of hGlyRS-S211F and hGlyRS-H418R in class IV multidendritic sensory neurons reduced their dendritic coverage by ∼35% and ∼46%, respectively (Figure 4C). Consistent with the results in motor neurons, simultaneous dGlyRS overexpression did not rescue the sensory neuron morphology defect. For hGlyRS-S211F, dGlyRS co-overexpression rather aggravated the sensory neuron morphology defect (Figure 4C).

Finally, we used THRONCAT, a recently developed method that allows for metabolic labelling of newly synthesized proteins with the non-canonical amino acid β-ethynylserine by the endogenous threonyl-tRNA synthetase (26) (see Materials and Methods for details). With this method, we observed that expression of the previously characterized hGlyRS-G240R variant in motor neurons reduced *de novo* protein synthesis by ∼19% (Supplementary Figure 2). Similarly, expression of hGlyRS-S211F or hGlyRS-H418R in motor neurons reduced protein synthesis by ∼17% and ∼18%, respectively (Figure 4D,E). Importantly, dGlyRS co-overexpression did not rescue this protein synthesis defect (Figure 4D,E). In all, these data show that the peripheral neuropathy phenotypes induced by hGlyRS-S211F and hGlyRS-H418R expression are not attributable to a dominant negative mechanism.

### Transgenic tRNA^Gly^ overexpression rescues peripheral neuropathy induced by hGlyRS-S211F and hGlyRS-H418R expression

We next investigated whether tRNA^Gly^ sequestration may be the pathogenic mechanism underlying PN-GlyRS caused by S211F and H418R mutations. To do so, we evaluated the effect of transgenic tRNA^Gly^ overexpression on peripheral neuropathy phenotypes and the mRNA translation defect induced by hGlyRS-S211F and hGlyRS-H418R. We used two previously described tRNA^Gly-GCC^ transgenic lines, a line that contains a BAC transgene with five tRNA^Gly-GCC^ genes that was inserted in the second and the third chromosome (overall resulting in 10 extra tRNA^Gly-GCC^ gene copies), and a tRNA^Gly-GCC^ ‘scramble’ transgenic line that contains 10 tRNA^Gly-GCC^ genes and results in higher level tRNA^Gly-GCC^ overexpression (∼13% overexpression for tRNA^Gly-GCC^ BAC and ∼30% for tRNA^Gly-GCC^ scramble)(12). Interestingly, the adult motor performance deficit induced by expression of hGlyRS-S211F or hGlyRS-H418R in motor neurons was rescued by both the tRNA^Gly-GCC^ BAC and tRNA^Gly-GCC^ scramble transgenes, and the degree of rescue was more pronounced for higher tRNA^Gly^ overexpression levels (Figure 5A,B). Consistent with this result, the reduced neuromuscular synapse length on larval muscle 8 induced by expression of hGlyRS-S211F or hGlyRS-H418R in motor neurons was significantly rescued by tRNA^Gly^ overexpression (Figure 5C). Furthermore, the sensory neuron dendritic morphology defect induced by hGlyRS-S211F or hGlyRS-H418R expression in class IV multidendritic sensory neurons was significantly rescued by tRNA^Gly^ overexpression (Figure 5D). Finally, THRONCAT revealed that the reduced protein synthesis following hGlyRS-S211F or hGlyRS-H418R expression in motor neurons is significantly rescued by tRNA^Gly^ overexpression (Figure 5E,F). Collectively, these data indicate that also for hGlyRS-S211F and hGlyRS-H418R mutations, tRNA^Gly^ sequestration is the underlying pathogenic mechanism.

**Figure 5:**
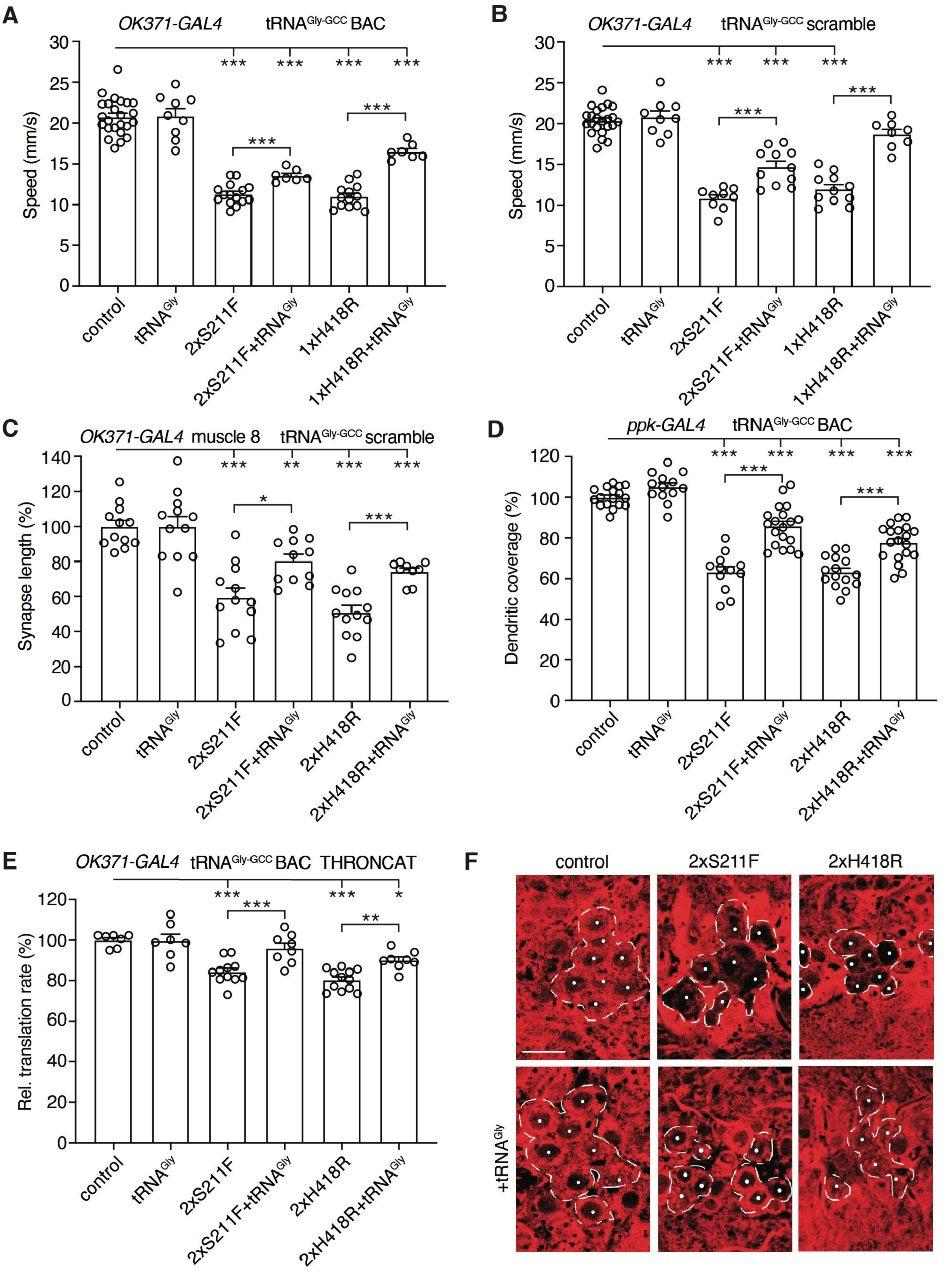
Transgenic tRNA^Gly^ overexpression rescues peripheral neuropathy phenotypes induced by hGlyRS-S211F and hGlyRS-H418R expression. A,B, Negative geotaxis climbing speed of 7-day-old female flies expressing hGlyRS-S211F or hGlyRS-H418R transgenes in motor neurons (*OK371-GAL4*), in the presence or absence of tRNA^Gly-GCC^ BAC (A) or tRNA^Gly-GCC^ scramble (B) transgenes. n = 7 to 24 (A) or 8 to 23 (B) groups of 10 flies per genotype; ***p<0.0005 by Brown-Forsythe and Welch ANOVA with Dunnett’s T3 multiple comparisons test (A) or one-way ANOVA with Sidak’s multiple comparisons test (B). **C,** Synapse length on distal muscle 8 of larvae expressing hGlyRS-S211F or hGlyRS-H418R transgenes in motor neurons (*OK371-GAL4*), in the presence or absence of tRNA^Gly-GCC^ overexpression. n = 9 to 12 animals per genotype; *p<0.05, **p<0.01, ***p<0.001 by Brown-Forsythe and Welch ANOVA with Dunnett’s T3 multiple comparisons test. **D,** Quantification of dendritic coverage (% of driver-only control) of class IV multidendritic sensory neurons in the larval body wall upon selective expression of hGlyRS-S211F or hGlyRS-H418R transgenes in these neurons (*ppk-GAL4*), in the presence or absence of tRNA^Gly-GCC^ overexpression. n = 12 to 19 animals per genotype; ***p<0.0001 by one-way ANOVA with Sidak’s multiple comparisons test. **E,F,** Quantification (E) and representative images (F) of relative translation rate (% of driver-only control) as determined by THRONCAT in motor neurons (*OK371-GAL4*) of larvae expressing hGlyRS-S211F or hGlyRS-H418R, in the presence or absence of tRNA^Gly-GCC^ overexpression. Motor neurons targeted by *OK371-GAL4* are delineated by a dashed line and nuclei of motor neurons are labeled by a white dot (F). n = 7 to 12 animals per genotype; *p<0.05, **p<0.01, ***p<0.001 by one-way ANOVA with Sidak’s multiple comparisons test. Scale bar: 10μm.

### Biochemical evidence for tRNA^Gly^ sequestration by hGlyRS S211F and H418R variants

To complement our *Drosophila* genetic data, we performed *in vitro* experiments to evaluate tRNA^Gly^ sequestration by hGlyRS S211F and H418R variants. We purified recombinant hGlyRS-WT, hGlyRS-S211F, and hGlyRS-H418R proteins and evaluated the effect of the mutations on hGlyRS dimer formation by size exclusion chromatography. Whereas hGlyRS-WT migrated predominantly as a dimer (dimer:monomer (D:M) ratio 9.1: 0.9), hGlyRS-S211F partitioned predominantly as monomer (D:M ratio 3.8: 6.2). For hGlyRS-H418R, the dimer was the predominant form but to a much lower fraction than hGlyRS-WT (D:M ratio 7.0: 3.0) (Figure 6A). We next used the hGlyRS dimer fraction to determine the association (*K_on_*) and release (*K_off_*) kinetic constants for tRNA^Gly-GCC^ to hGlyRS-WT, hGlyRS-S211F and hGlyRS-H418R. Both hGlyRS-S211F and hGlyRS-H418R variants were able to bind tRNA^Gly^, albeit with a reduced affinity (∼3-fold and ∼5-fold fold lower *K_on_* values than WT, respectively, Figure 6B). Importantly, these variants displayed markedly slower tRNA^Gly^ release kinetics (*K_off_*), with more than 90% of traces showing no tRNA^Gly^ release within the experimental time frame (Figure 6B). These data are consistent with tRNA^Gly^ sequestration by hGlyRS-S211F and hGlyRS-H418R variants.

**Figure 6:**
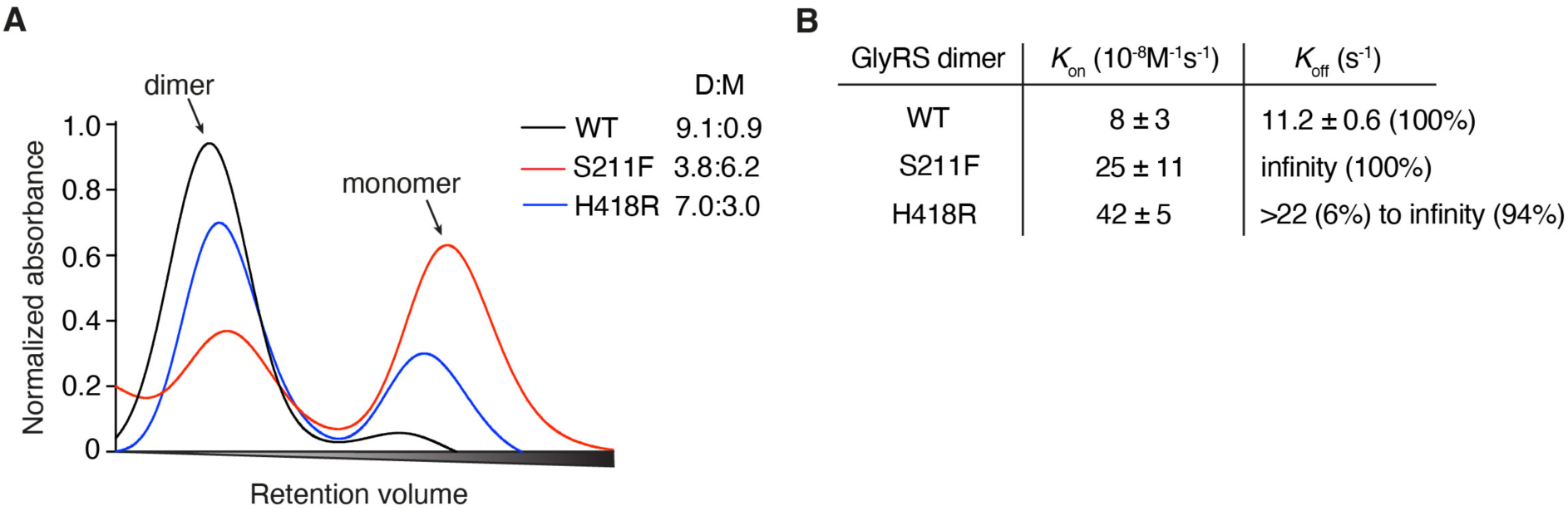
The S211F and H418R mutations alter GlyRS dimerization and the release kinetics of the cognate tRNA^Gly^. A, Size-exclusion chromatography of purified recombinant hGlyRS proteins. D:M, dimer-to-monomer ratio. **B,** *K_on_* and *K_off_* values of tRNA^Gly-GCC^ binding and release, respectively, to the dimer form of the indicated hGlyRS variants. The percentage in parentheses denotes the frequency of a measured value.

## DISCUSSION

For many genetic diseases, multiple disease-causing mutations in the same gene have been identified, often distributed throughout the coding region and spread across diverse functional domains. For the vast majority of these diseases, it is unclear whether all mutations cause disease through the same molecular mechanism. Here, we addressed this question for peripheral neuropathy associated with mutations in *GARS1*, encoding GlyRS. Previously, we provided genetic and biochemical evidence for seven PN-GlyRS mutations, that sequestration of tRNA^Gly^ and consequent depletion of the cellular tRNA^Gly^ pool is the underlying molecular mechanism (12). The insufficient availability of tRNA^Gly^ substrate for aminoacylation by WT GlyRS, derived from the WT *GARS1* allele in heterozygous patients, leads to insufficient glycyl-tRNA^Gly^ to be used by the translating ribosome. This in turn triggers ribosome stalling at glycine codons (12) and activation of the ISR through the eIF2α kinase GCN2 (11). This mechanism also provides an explanation for the protein synthesis defect observed in PN-GlyRS *Drosophila* and mouse models (10,11).

Here, we evaluated whether all known PN-GlyRS variants cause disease through tRNA^Gly^ sequestration, or alternatively, whether a subset of these mutations may trigger peripheral neuropathy through a dominant negative effect on GlyRS aminoacylation activity. Indeed, because GlyRS functions as a homodimer with a single tRNA^Gly^ interacting with both GlyRS monomers (38), a GlyRS dimer consisting of one WT and one mutant monomer may fail to aminoacylate tRNA^Gly^, provided that the mutation results in loss of aminoacylation activity but does not inhibit dimerization with WT GlyRS. In such a dominant negative scenario, the aminoacylation-active GlyRS population, which consists of WT GlyRS homodimers, may be reduced by 75%. In turn, 75% reduction of tRNA^Gly^ aminoacylation capacity may substantially lower the levels of glycyl-tRNA^Gly^ available for translation, thus potentially triggering peripheral neuropathy. In *Drosophila* PN-GlyRS models, a dominant negative mechanism would require dimerization between endogenous dGlyRS and transgenic hGlyRS, which we previously demonstrated to occur (10). The plausibility of a dominant-negative mechanism underlying PN-GlyRS was suggested by our finding that reducing endogenous dGlyRS levels by transgenic RNAi-mediated knock-down is sufficient to induce peripheral neuropathy phenotypes and to substantially inhibit *de novo* protein synthesis (Figure 1).

To evaluate a possible dominant negative mechanism and because all PN-GlyRS mutations in *Drosophila* and mouse models generated thus far increase the net positive charge of the GlyRS protein (10,16,39–41), we decided to generate three novel *Drosophila* PN-GlyRS models for two mutations that do not alter GlyRS protein charge (S211F and H418R)(32,34), and the single GlyRS mutation described that reduces positive charge (K456Q)(35,37). The S211F and H418R mutations were previously reported to result in severely reduced tRNA^Gly^ aminoacylation activity (29,33), whereas the effect of the K456Q mutation on aminoacylation activity has not been reported. Very similar to our previously reported *Drosophila* PN-GlyRS models (10), expression of hGlyRS-S211F or hGlyRS-H418R in motor and sensory neurons induced peripheral neuropathy phenotypes and substantially inhibited *de novo* protein synthesis in motor neurons (Figure 4, 5). This phenotypic similarity supports the notion of a similar underlying pathogenic mechanism.

To evaluate whether either a dominant negative mechanism or tRNA^Gly^ sequestration may underlie PN-GlyRS caused by S211F and H418R mutations, we studied the effect of transgenic overexpression of WT dGlyRS or tRNA^Gly^, respectively. For both hGlyRS-S211F and hGlyRS-H418R variants, dGlyRS overexpression did not rescue peripheral neuropathy or inhibition of protein synthesis, providing a strong argument against a dominant negative mechanism (Figure 4). This result is consistent with the finding that transgenic overexpression of WT hGlyRS did not rescue peripheral neuropathy phenotypes in two PN-GlyRS mouse models (17). In contrast, tRNA^Gly^ overexpression dosage-dependently rescued both peripheral neuropathy phenotypes and protein synthesis (Figure 5), providing genetic evidence for depletion of the cellular tRNA^Gly^ pool due to tRNA^Gly^ sequestration as the underlying pathogenic mechanism. In addition, we provide biochemical evidence of tRNA^Gly^ sequestration by hGlyRS-S211F and hGlyRS-H418R variants (Figure 6). Thus, addition of positive charge is not required for a PN-GlyRS mutation to induce tRNA^Gly^ sequestration. This is consistent with our previous finding that the hGlyRS-L129P and hGlyRS-E279D mutations, which do not alter protein charge, clearly induce tRNA^Gly^ sequestration in biochemical experiments (12).

For our previously characterized PN-GlyRS *Drosophila* models (E71G, G240R, G526R mutations) (10), we had shown that tRNA^Gly^ overexpression at least partially rescues peripheral neuropathy phenotypes and the protein synthesis defect (12).

Here, we evaluated the effect of dGlyRS overexpression on these phenotypes. For hGlyRS-E71G, dGlyRS overexpression did not rescue peripheral neuropathy or protein synthesis (Figure 2), arguing against a dominant negative mechanism. Given that the E71G mutation does not significantly affect tRNA^Gly^ aminoacylation activity (10,29,30) or GlyRS dimerization (12), this result could be expected. In contrast, G240R and G526R severely reduce aminoacylation activity (10,30,31,33), which may hint to a dominant negative mechanism. Interestingly, dGlyRS overexpression partially rescued peripheral neuropathy and protein synthesis defects induced by the G240R and G526R variants. A possible explanation for this result is that in addition to tRNA^Gly^ sequestration, these two GlyRS variants also induce a dominant negative effect, which may, at least in part, contribute to the insufficient glycyl-tRNA^Gly^ production and supply to the ribosome. An alternative explanation for this observation is that WT and mutant GlyRS species may compete for binding to the limited cellular tRNA^Gly^ pool. In this scenario, a substantial increase in dGlyRS levels may increase the tRNA^Gly^ fraction interacting with and aminoacylated by WT dGlyRS, thus increasing glycyl-tRNA^Gly^ levels available to the translating ribosome. Consistent with this hypothesis, the fact that the G240R mutation was reported to substantially reduce hGlyRS dimerization capacity (30) argues against a dominant negative mechanism. Co-immunoprecipitation experiments in transfected cells previously suggested that hGlyRS-G526R may retain or even strengthen GlyRS dimerization capacity (30,31), but size-exclusion chromatography revealed that recombinant hGlyRS-G526R protein predominantly occurs as a monomer (12).

Although our data show a clear correlation between the degree to which PN-GlyRS variants inhibit global protein synthesis and the severity of peripheral neuropathy phenotypes, this correlation is not perfect. Furthermore, while tRNA^Gly^ overexpression substantially rescues peripheral neuropathy phenotypes of all PN-GlyRS *Drosophila* models, the rescue is not complete. These observations may be explained by the fact that the UAS-hGlyRS transgenes allow expression of both the cytosolic and the mitochondrial hGlyRS isoforms, depending on the translation start site used. Because in higher eukaryotes, mitochondrial tRNAs are encoded by the mitochondrial genome and mitochondria typically do not import nuclear encoded tRNA (42), mitochondrial translation defects and toxicity induced by PN-GlyRS variants are likely not rescued by tRNA^Gly^ overexpression. Moreover, additional pathogenic mechanisms, such as axonal transport defects induced by decreased α-tubulin acetylation (43,44), may not be fully rescued by tRNA^Gly^ overexpression. In spite of the possible contribution of mitochondrial toxicity induced by PN-GlyRS variants (45), the cytosolic toxicity induced by tRNA^Gly^ sequestration is likely a critical contributor to peripheral neuropathy phenotypes, because (i) in PN-GlyRS mouse models, transgenic tRNA^Gly^ overexpression fully rescues peripheral neuropathy phenotypes (12), and (ii) all of the seven other PN-associated aaRS genes exclusively encode the cytosolic aaRS, with the corresponding mitochondrial aaRSs being encoded by different genes.

Size-exclusion chromatography revealed that GlyRS-S211F predominantly occurs as a monomer, similar to some other PN-GlyRS variants (e.g. L129P, C157R, G240R) (12,30). We previously reported that PN-GlyRS monomers are able to bind and sequester tRNA^Gly^ (12), which we anticipate may also be the case for GlyRS-S211F. Consistent with this notion, peripheral neuropathy phenotypes induced by GlyRS S211F, C157R and G240R variants in *Drosophila* and mouse models are rescued by transgenic tRNA^Gly^ overexpression.

In sharp contrast to hGlyRS-S211F and hGlyRS-H418R, expression of hGlyRS-K456Q induced neither peripheral neuropathy phenotypes, nor inhibition of protein synthesis. Although the K456Q variant was previously classified as likely pathogenic (35), according to the most recent American College of Medical Genetics and Genomics (ACMG) guidelines (46), this variant would be classified as a variant of uncertain significance. Our data in *Drosophila* (Figure 3) indicate that this hGlyRS variant may not be pathogenic. Because the K456Q variant is the only one of the 35 reported PN-GlyRS variants that eliminates a positive charge from the GlyRS protein, our data further suggest that mutations that render the overall GlyRS charge more negative than the WT protein may not be compatible with PN-GlyRS pathogenicity in heterozygous patients. In contrast, 4 of the 7 *GARS1* mutations reported in compound heterozygous patients with biallelic *GARS1* mutations remove a positive charge from the GlyRS protein (1). These patients display systemic mitochondrial disease or multisystem developmental syndrome (45,47–50), and these phenotypes are likely attributable to a substantial reduction of tRNA^Gly^ aminoacylation activity (1). The systemic mitochondrial disease features suggest that mitochondrial mRNA translation may be predominantly affected, leading to mitochondrial respiratory chain dysfunction (45,47,48,50).

Overall, our data indicate that it is highly likely that all PN-GlyRS mutations trigger peripheral neuropathy through a common tRNA^Gly^ sequestration mechanism. From a therapeutic perspective, this suggests that elevating tRNA^Gly^ levels in the affected motor and sensory neurons may constitute a viable therapeutic approach for all PN-GlyRS patients, irrespective of the disease-causing mutation.

## DATA AVAILABILITY

All data are presented in the main manuscript and supporting files. Raw data and images underlying this article will be shared on request to the corresponding authors.

## AUTHOR CONTRIBUTIONS

N.M., E.F.J.S., V.L., M.P.M., M.M., P.v.L., C.S., and N.v.B. performed experiments. All authors analyzed and discussed the data. ES conceived, initiated, and supervised the project. ES wrote the manuscript with contribution of N.M. and E.F.J.S. All authors contributed to the experimental design and interpretation and commented on the manuscript.

## ACKNOWLEDGEMENTS

We thank Robin Thompson for generating the plasmids for purification of the hGlyRS-S211F and hGlyRS-H418R variants, Bob Ignacio and Kimberly Bonger for providing βES for THRONCAT experiments, and the General Instruments Facility (Faculty of Science, Radboud University) for advice on image acquisition and analysis.

---

**Supplementary Figure 1:**
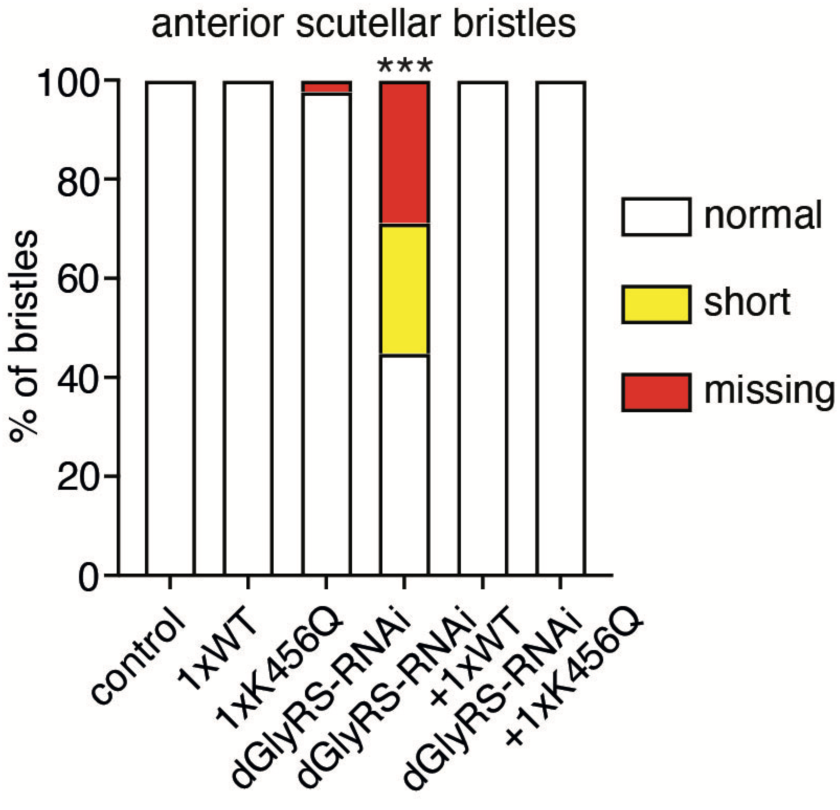
Quantification of the percentage of normal (long), short, or missing anterior scutellar bristles in the presence or absence of dGlyRS knock-down, and with or without simultaneous co-expression of hGlyRS-WT or hGlyRS-K456Q. n = 36 to 118 bristles per genotype; ***p<0.0001 by Fisher’s exact test.

**Supplementary Figure 2:**
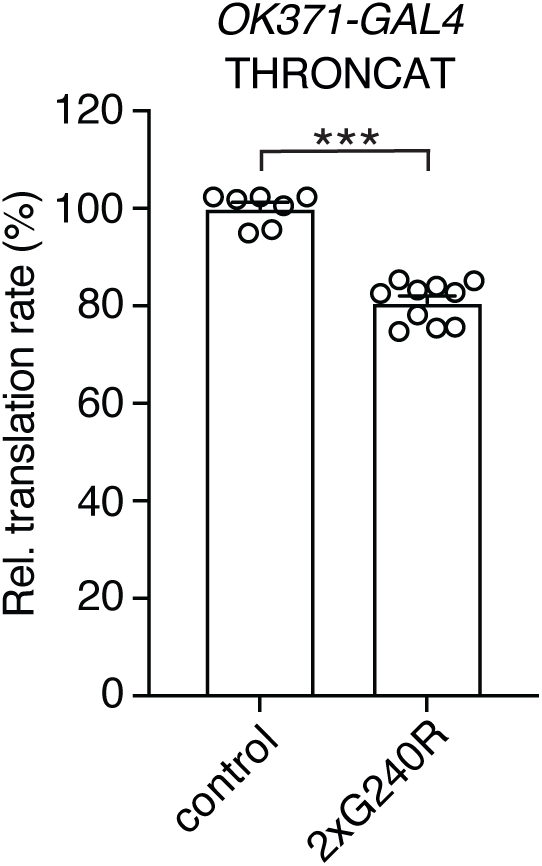
Relative translation rate (% of driver-only control) as determined by THRONCAT in motor neurons (*OK371-GAL4*) of larvae expressing hGlyRS-G240R. n = 7 to 12 animals per genotype; ***p<0.0001 by Mann-Whitney test. The control data points are the same as in Figure 5E, because the 2xG240R condition was included in the same larger experiment.

**Supplementary Table 1:**
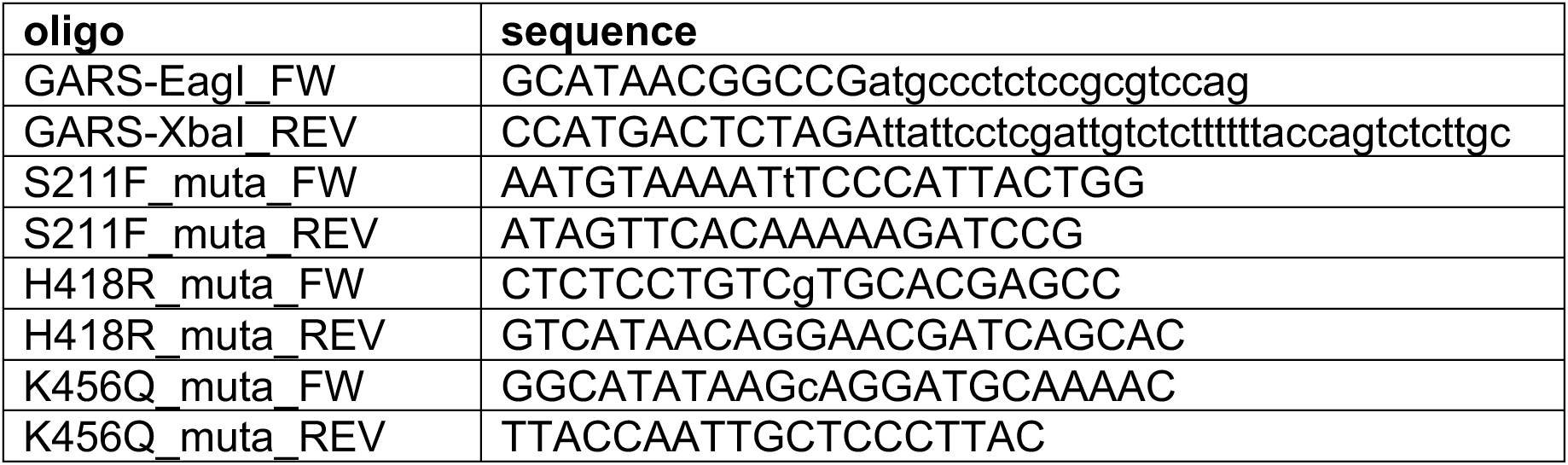
Sequences of primers used for cloning and mutagenesis of human GARS1 cDNA.

**Supplementary Table 2:**
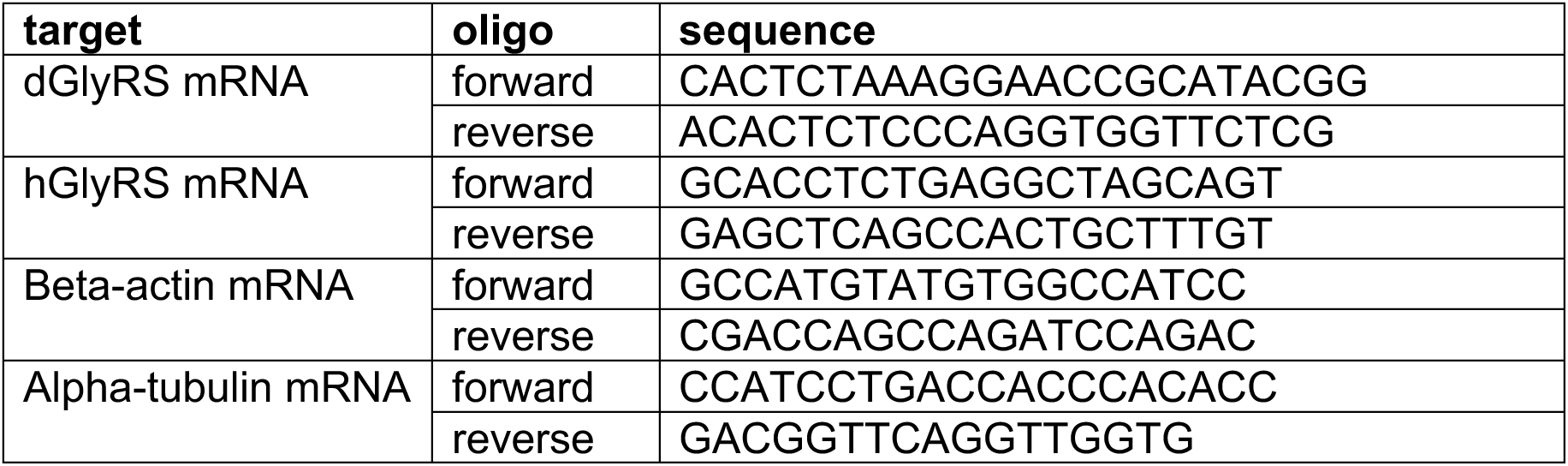
Sequences of primers used for quantitative real-time PCR (qPCR).

**Supplementary Table 3:** Characteristics of PN-GlyRS variants used in this study. The effect of PN-GlyRS variants on net GlyRS protein charge, aminoacylation activity, evolutionary conservation of the affected amino acid residues, and genetic evidence to support disease causality are shown. Segregation of the mutation with disease in a family provides stronger genetic evidence than mutations found in single patients. Furthermore, a summary of peripheral neuropathy-like phenotypes in *Drosophila* PN- GlyRS models is provided, including the effect of expression of PN-GlyRS variants on protein synthesis in motor neurons, developmental lethality induced by ubiquitous expression, impaired adult climbing behavior induced by motor neuron (*OK371-GAL4*) expression as evaluated in the negative geotaxis assay, reduced dendritic coverage of class IV multidendritic sensory neurons upon selective expression in these neurons (*ppk-GAL4*). Symbols: = not altered; ↓, mild reduction; ↓,↓, moderate reduction; ↓,↓,↓, severe reduction; - no phenotype; + mild phenotype; ++ moderate phenotype; +++ severe phenotype. AA: amino acid; NA: not applicable; ND: not determined.

| GlyRS mutation | effect on net GlyRS charge | evolutionary conservation of mutated AA residue | segregation with disease in a family | aminoacylation activity | protein synthesis | developmental lethality induced by ubiquitous expression | impaired climbing behavior | reduced dendritic coverage of multidendritic sensory neurons | references |
| --- | --- | --- | --- | --- | --- | --- | --- | --- | --- |
| WT | = | NA | NA | = | = | - | - | - |  |
| E71G | +1 | yeast | yes | = | ↓↓ | ++ | ++ | + | (5,10,29,30) |
| S211F | = | C. elegans | yes | ↓↓↓ | ↓↓↓ | +++ | ++ | ++ | (32,33), this study |
| G240R | +1 | Drosophila | yes | ↓↓ | ↓↓↓ | +++ | +++ | ++ | (5,10,29,30,33) |
| H418R | = | yeast | yes | ↓↓ | ↓↓↓ | +++ | ++ | ++ | (29,33,34), this study |
| K456Q | -1 | yeast | no | = | = | - | - | - | (35), this study |
| G526R | +1 | yeast | yes | ↓↓↓ | ↓↓↓ | +++ | +++ | ++ | (5,10,29,30,33, 36) |

